# Sequence action representations contextualize during early skill learning

**DOI:** 10.1101/2024.08.15.608189

**Authors:** Debadatta Dash, Fumiaki Iwane, William Hayward, Roberto Salamanca-Giron, Marlene Bonstrup, Ethan R. Buch, Leonardo G. Cohen

## Abstract

Activities of daily living rely on our ability to acquire new motor skills composed of precise action sequences. Here, we asked if the millisecond-level neural representation of an action performed at different contextual sequence locations within a skill differentiates or remains stable during early motor learning. We first optimized machine learning decoders predictive of sequence-embedded finger movements from magnetoencephalographic (MEG) activity. Using this approach, we found that the neural representation of the same action performed in different contextual sequence locations, progressively differentiated—primarily during rest intervals of early learning (offline)—correlating with skill gains. In contrast, representational differentiation during practice (online) did not reflect learning. The regions contributing to this representational differentiation evolved with learning, shifting from the contralateral pre- and post-central cortex during early learning (trials 1–11) to increased involvement of the superior and middle frontal cortex once skill performance plateaued (trials 12–36). Thus, the neural substrates supporting finger movements and their representational differentiation during early skill learning differ from those supporting stable performance during the subsequent skill plateau period. Representational contextualization extended to Day 2, exhibiting specificity for the practiced skill sequence. Altogether, our findings indicate that sequence action representations contextually differentiate during early skill learning, an issue relevant to brain-computer interface applications in neurorehabilitation.

## Introduction

Motor learning is required to perform a wide array of activities of daily living, intricate athletic endeavors, and professional skills. Whether it’s learning to type more quickly on a keyboard [1], improve one’s tennis game [2], or play a piece of music on the piano [3]– all these skills require the ability to execute sequences of actions with precise temporal coordination. Action sequences thus form the building blocks of fine motor skills [4]. Practicing a new motor skill elicits rapid performance improvements (early learning) [1] that precede skill performance plateaus [5]. Skill gains during early learning accumulate over rest periods (micro-offline) interspersed with practice [1, 6–10], and are up to four times larger than offline performance improvements reported following overnight sleep [1]. During this initial interval of prominent learning, retroactive interference immediately following each practice interval reduces learning rates relative to interference after passage of time, consistent with stabilization of the motor memory [11]. Micro-offline gains observed during early learning are reproducible [7, 10–13] and are similar in magnitude even when practice periods are reduced by half to 5 seconds in length, thereby confirming that they are not merely a result of recovery from performance fatigue [11]. Additionally, they are unaffected by the random termination of practice periods, which eliminates the possibility of predictive motor slowing as a contributing factor [11]. Collectively, these behavioral findings point towards the interpretation that micro-offline gains during early learning represent a form of memory consolidation [1].

This interpretation has been further supported by brain imaging and electrophysiological studies linking known memory-related networks and consolidation mechanisms to rapid offline performance improvements. In humans, the rate of hippocampo-neocortical neural replay predicts micro-offline gains [6]. Consistent with these findings, Chen et al. [12] and Sjøgård et al. [13] furnished direct evidence from intracranial human EEG studies, demonstrating a connection between the density of hippocampal sharp-wave ripples (80-120 Hz)—recognized markers of neural replay—and micro-offline gains during early learning. Further, Griffin et al. reported that neural replay of task-related ensembles in the motor cortex of macaques during brief rest periods— akin to those observed in humans [1, 6–8, 14]—are not merely correlated with, but are causal drivers of micro-offline learning [15]. Specifically, the same reach directions that were replayed the most during rest breaks showed the greatest reduction in path length (i.e. – more efficient movement path between two locations in the reach sequence) during subsequent trials, while stimulation applied during rest intervals preceding performance plateau reduced reactivation rates and virtually abolished micro-offline gains [15]. Thus, converging evidence in humans and non-human primates across indirect non-invasive and direct invasive recording techniques link hippocampal activity, neural replay dynamics and offline skill gains in early motor learning that precede performance plateau.

During skill learning, the neural representation of a sequential skill binds discrete individual actions (e.g. - single piano keypress) into complex, temporally and spatially precise sequence representations (e.g. - a refrain from a piece of music) [16–20]. After a skill is learned over extended periods (i.e., weeks), the neural representation of the sequence changes significantly [20], while the representation of its individual action components (e.g., finger movements) does not [21]. On the other hand, it is not known whether individual sequence action representations differentiate or remain stable during the early stages of skill learning, when the memory is still not fully formed [1]. Furthermore, it is unknown whether the neural representations of identical movements, performed at different positions within a skill sequence (i.e. - the skill *context*), differentiate with learning—an important consideration for advancing robust brain-computer interface (BCI) applications [22–26].

Examining the millisecond-level differentiation of discrete action representations during learning is challenging, as evolving neural dynamics concurrently encode skill sequences and their individual action components [20, 27] across multiple spatial scales [28]. To address this problem, we first optimized a multi-scale decoder aimed at predicting keypress actions from magnetoencephalographic (MEG) neural activity. Using this optimized approach, we report that an individual sequence action representation differentiates depending on the sequence context and correlates with early skill learning. This representational contextualization developed predominantly over rest rather than during practice intervals—in parallel with rapid consolidation of skill.

## Results

Participants engaged in a well characterized sequential skill learning task. [1, 6, 11] that involved repetitive typing of a sequence (4-1-3-2-4) performed with their (non-dominant) left hand over 36 trials with alternating periods of 10s *practice* and 10s *rest* (*inter-practice rest*; *Day 1 Training*; **Figure 1A**), a practice schedule that minimizes reactive inhibition effects [11, 29] (see **Methods**). Individual keypress times and finger keypress identities were recorded and used to quantify skill as the correct sequence speed (keypresses/s) [1].

**Figure 1.**
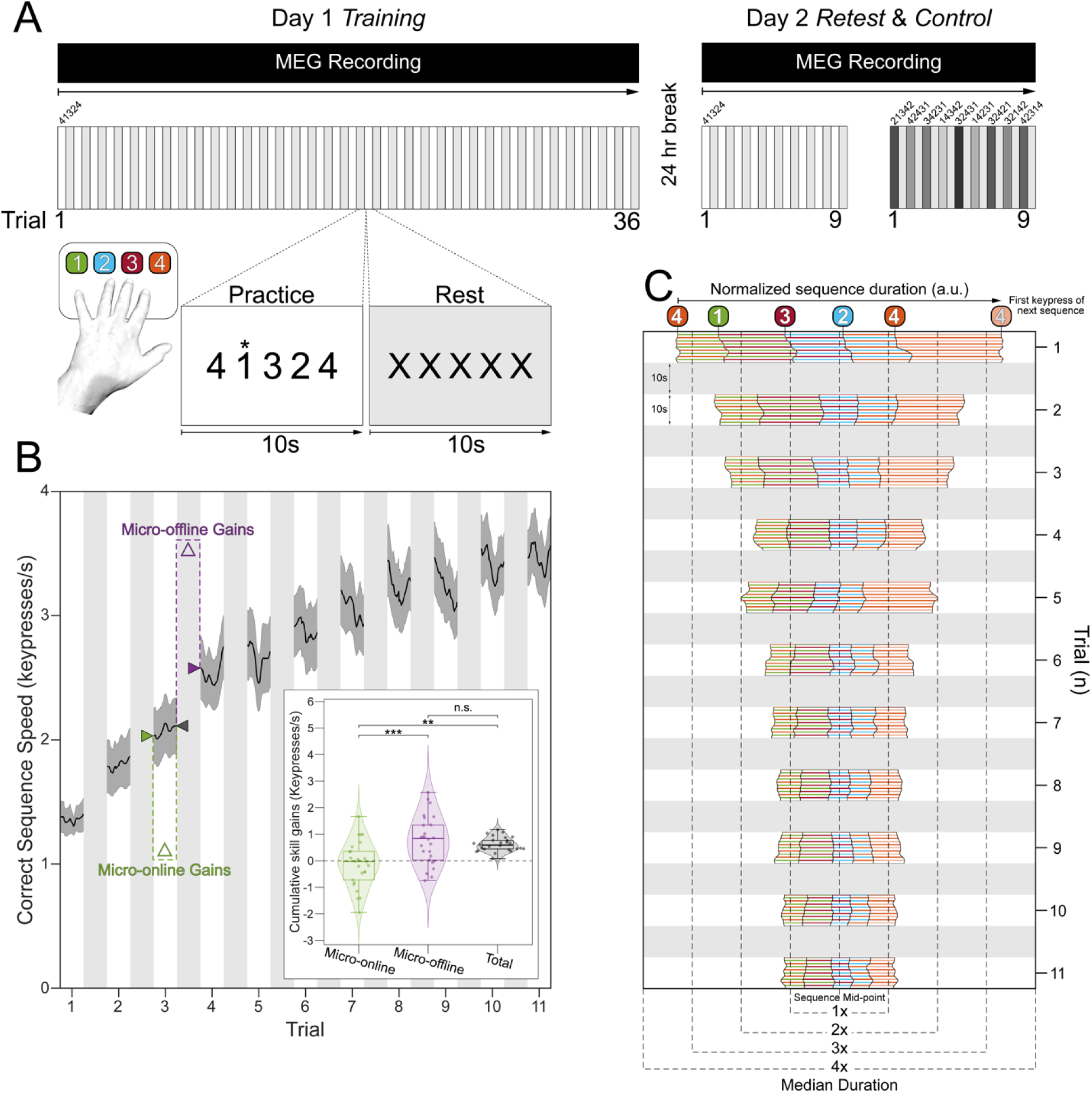
Experimental design and behavioral performance. *A. Skill learning task.* Participants engaged in a procedural motor skill learning task, which required them to repeatedly type a keypress sequence, “4 − 1 − 3 − 2 − 4” (1 = little finger, 2 = ring finger, 3 = middle finger, and 4 = index finger) with their non-dominant, left hand. The *Day 1 Training* session included 36 trials, with each trial consisting of alternating 10s practice and rest intervals. The rationale for this task design was to minimize reactive inhibition effects during the period of steep performance improvements (early learning) [11, 29] (see **Methods**). After a 24-hour break, participants were retested on performance of the same sequence (4-1-3-2-4) for 9 trials (*Day 2 Retest)* to inform on the generalizability of the findings over time and MEG recording sessions, as well as single-trial performance on 9 different control sequences (*Day 2 Control*; 2-1-3-4-2, 4-2-4-3-1, 3-4-2-3-1, 1-4-3-4-2, 3-2-4-3-1, 1-4-2-3-1, 3-2-4-2-1, 3-2-1-4-2, and 4-2-3-1-4) to inform on specificity of the findings to the learned skill. MEG was recorded during both Day 1 and Day 2 sessions with a 275-channel CTF magnetoencephalography (MEG) system (CTF Systems, Inc., Canada). ***B. Skill Learning***. As reported previously^1^, participants on average reached 95% of peak performance by trial 11 of the *Day 1 Training* session (see **Figure 1 – figure supplement 1A** for results over all *Day 1 Training* and *Day 2 Retest* trials). At the group level, total early learning was exclusively accounted for by micro-offline gains during inter-practice rest intervals (Figure 1B**, inset**; F [2,75] = 14.79, *p* = 3.86x10^-6^; micro-online vs. micro-offline: *p* = 7.98x10^-6^; micro-online vs. total: *p* = 0.0002; micro-offline vs. total: *p* = 0.669). These results were not impacted by potential preplanning effects on initial skill performance [30] since alternative measurements of cumulative micro-online and - offline gains remain unchanged after omission of the first 3 keypresses in each trial from the correct sequence speed computation (paired t-tests; micro-online: *t_25_* = -0.0223, *p* = 0.982; micro-offline: *t_25_* = -0.879, *p* = 0.388). **C. Keypress transition time (KTT) variability.** Distribution of KTTs normalized to the median correct sequence time for each participant and centered on the mid-point for each full sequence iteration during early learning (see **Figure 1 – figure supplement 1B** for results over all *Day 1 Training* and *Day 2 Retest* trials). Note the initial variability of the relative KTT composition of the sequence (i.e., – 4-1, 1-3, 3-2, 2-4, 4-4), before it stabilizes in the early learning period.

Participants reached 95% of maximal skill (i.e., - Early Learning) within the initial 11 practice trials (**Figure 1B)**, with improvements developing over inter-practice rest periods (micro-offline gains) accounting for almost all total learning across participants (**Figure 1B, inset)**[1]. In addition to the reduction in sequence duration during early learning, individual keypress transition times became more consistent across repeated sequence iterations (**Figure 1C**). On average across subjects, 2.32% ± 1.48% (mean ± SD) of all keypresses performed were errors, which were evenly distributed across the four possible keypress responses. While errors increased progressively over practice trials, they did so in proportion to the increase in correct keypresses, so that the overall ratio of correct-to-incorrect keypresses remained stable over the training session.

On the following day, participants were retested on performance of the same sequence (4-1-3-2-4) over 9 trials (*Day 2 Retest*), as well as on the single-trial performance of 9 different untrained control sequences (*Day 2 Controls*: 2-1-3-4-2, 4-2-4-3-1, 3-4-2-3-1, 1-4-3-4-2, 3-2-4-3-1, 1-4-2-3-1, 3-2-4-2-1, 3-2-1-4-2, and 4-2-3-1-4). As expected, an upward shift in performance of the trained sequence (0.68 ± SD 0.56 keypresses/s; *t* = 7.21, *p* < 0.001) was observed during *Day 2 Retest*, indicative of an overnight skill consolidation effect (**Figure 1 – figure supplement 1A**).

### Keypress actions are represented in multi-scale hybrid-space manifolds

We investigated the differentiation of neural representations of the same index finger keypress performed at different positions of the skill sequence. A set of decoders were constructed to predict keypress actions from MEG activity as a function of both the learning state and the ordinal position of the keypress within the sequence. We first characterized the spectral and spatial features of keypress state representations by comparing performance of decoders constructed around broadband (1-100Hz) or narrowband [delta-(1-3Hz), theta- (4-7Hz), alpha- (8-14 Hz), beta- (15-24 Hz), gamma- (25-50Hz) and high gamma-band (51-100Hz)] MEG oscillatory activity. We found that decoders trained on broadband activity consistently outperformed those trained on narrowband activity. Whole-brain parcel-space (70.11% ± SD 7.11% accuracy; n = 148 brain regions; *t* = 1.89, *p* = 0.035, *df* = 25, Cohen’s *d* = 0.17, **Figure 2A**; also see **Figure 2B** for topographic map of feature importance scores) and voxel-space (74.51% ± SD 7.34% accuracy; n = 15684; *t* = 7.18, *p* < 0.001, *df* = 25, Cohen’s *d* = 0.76, **Figure 2A**; also see **Figure 2C** for topographic map of feature importance scores) [31] decoders exhibited greater accuracy than all regional voxel-space decoders constructed from individual brain areas (**Figure 2D**; maximum accuracy of 68.77%± SD 7.6%; see also **Figure 2 – figure supplements 1 & 2)**. Thus, optimal decoding required information from multiple brain regions, predominantly contralateral to the hand engaged in the skill task (**Figure 2B** and **C**).

**Figure 2.**
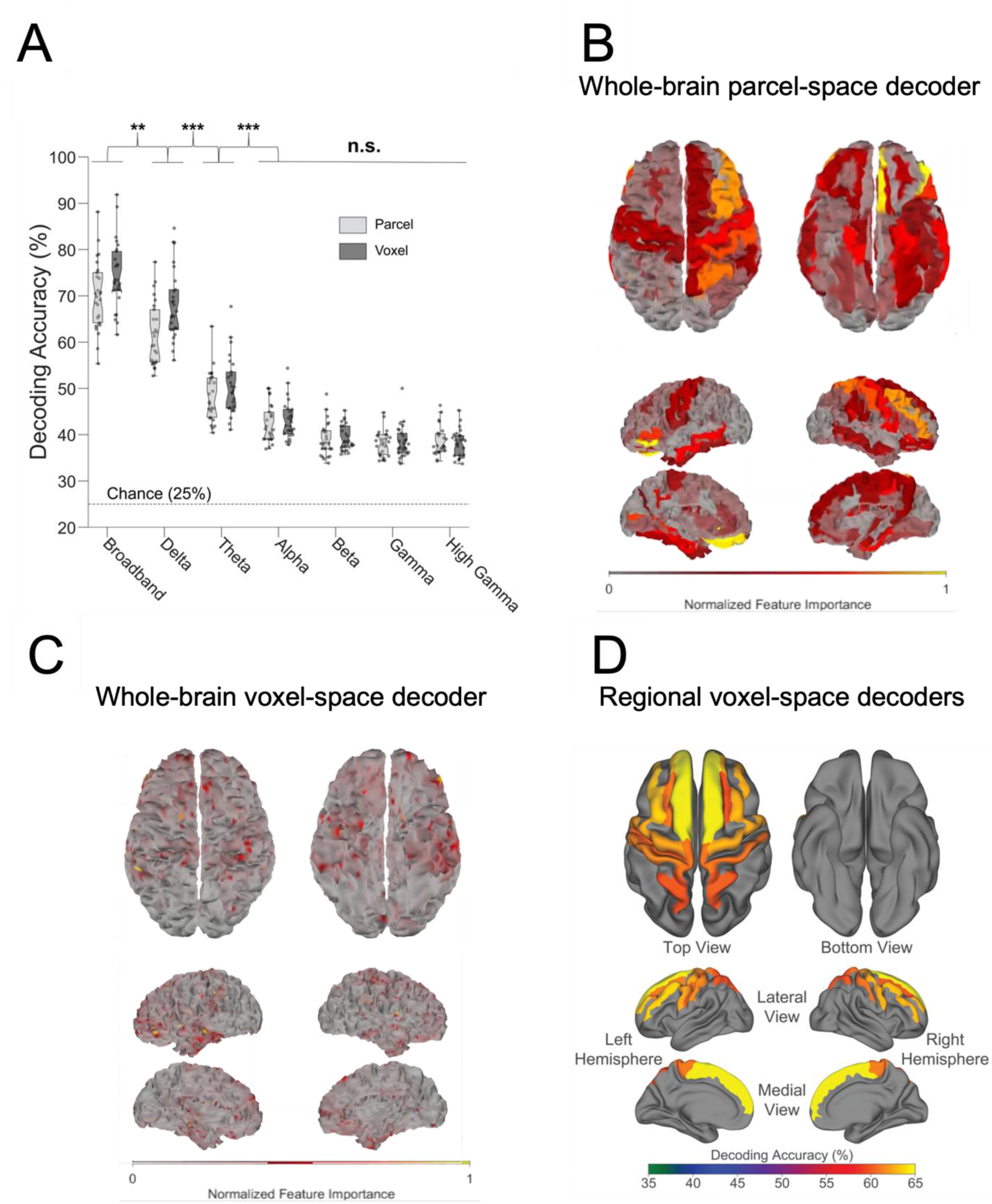
Spatial and oscillatory contributions to neural decoding of finger identities. *A) Contribution of whole-brain oscillatory frequencies to decoding.* When trained on broadband activity relative to narrow frequency band features, decoding accuracy (i.e. - test sample performance) was highest for whole-brain voxel-space (74.51% ± SD 7.34%, *t* = 8.08, *p* < 0.001) and parcel-space (70.11% ± SD 7.11%, *t* = 13.22, *p* < 0.001) MEG activity. Thus, decoders trained on whole-brain broadband data consistently outperformed those trained on narrowband activity. Dots depict decoding accuracy for each participant. \**p* < 0.05, \*\**p*< 0.01, \*\*\**p*< 0.001, ns.: not significant. *B) Whole-brain parcel-space decoding.* Color-coded standard (FreeSurfer *fsaverage*) brain surface plot displaying the relative importance of individual brain regions (parcels) to broadband whole-brain parcel-space decoding performance (far-left light gray box plot in *A*). *C) Whole-brain voxel space decoding.* Color-coded standard (FreeSurfer *fsaverage*) brain surface plot displaying the relative importance of individual voxels to broadband whole-brain voxel-space decoding performance (far-left dark gray box plot in *A*). *D) Regional voxel-space decoding.* Broadband voxel-space decoding performance for top-ranked brain regions across the group is displayed on a standard (FreeSurfer *fsaverage*) brain surface and color-coded by accuracy. Note that while whole-brain parcel- and voxel-space decoders relied more on information from brain regions contralateral to the engaged hand, regional voxel-space decoders performed similarly for bilateral sensorimotor regions.

Next, given that the brain simultaneously processes information more efficiently across multiple spatial and temporal scales [28, 32, 33], we asked if the combination of lower resolution whole-brain and higher resolution regional brain activity patterns further improve keypress prediction accuracy. We constructed hybrid-space decoders (N = 1295 ± 20 features; **Figure 3A)** combining whole-brain parcel-space activity (n = 148 features; **Figure 2B**) with regional voxel-space activity from a data-driven subset of brain areas (n = 1147 ± 20 features; **Figure 2D**). This subset covers brain regions showing the highest regional voxel-space decoding performances (top regions across all subjects shown in **Figure 2D; Methods *– Hybrid Spatial Approach***). Accuracy was higher for hybrid- (78.15% ± SD 7.03%; weighted mean F1 score of 0.78 ± SD 0.07) than for voxel- (74.51% ± SD 7.34%; paired *t*-test: *t* = 6.30, *p* < 0.001, *df* = 25, Cohen’s *d* = 0.39) and parcel-space decoders (70.11% ± SD 7.48%; paired *t*-test: *t* = 12.08, *p* < 0.001, *df* = 25, Cohen’s *d* = 0.98, **Figure 3B, Figure 3 – figure supplement 1,6**). Note that while features from contralateral brain regions were more important for whole-brain decoding (in both parcel- and voxel-spaces), regional voxel-space decoders performed best for bilateral sensorimotor areas on average across the group. Thus, a multi-scale hybrid-space representation best characterizes the keypress action manifolds.

**Figure 3.**
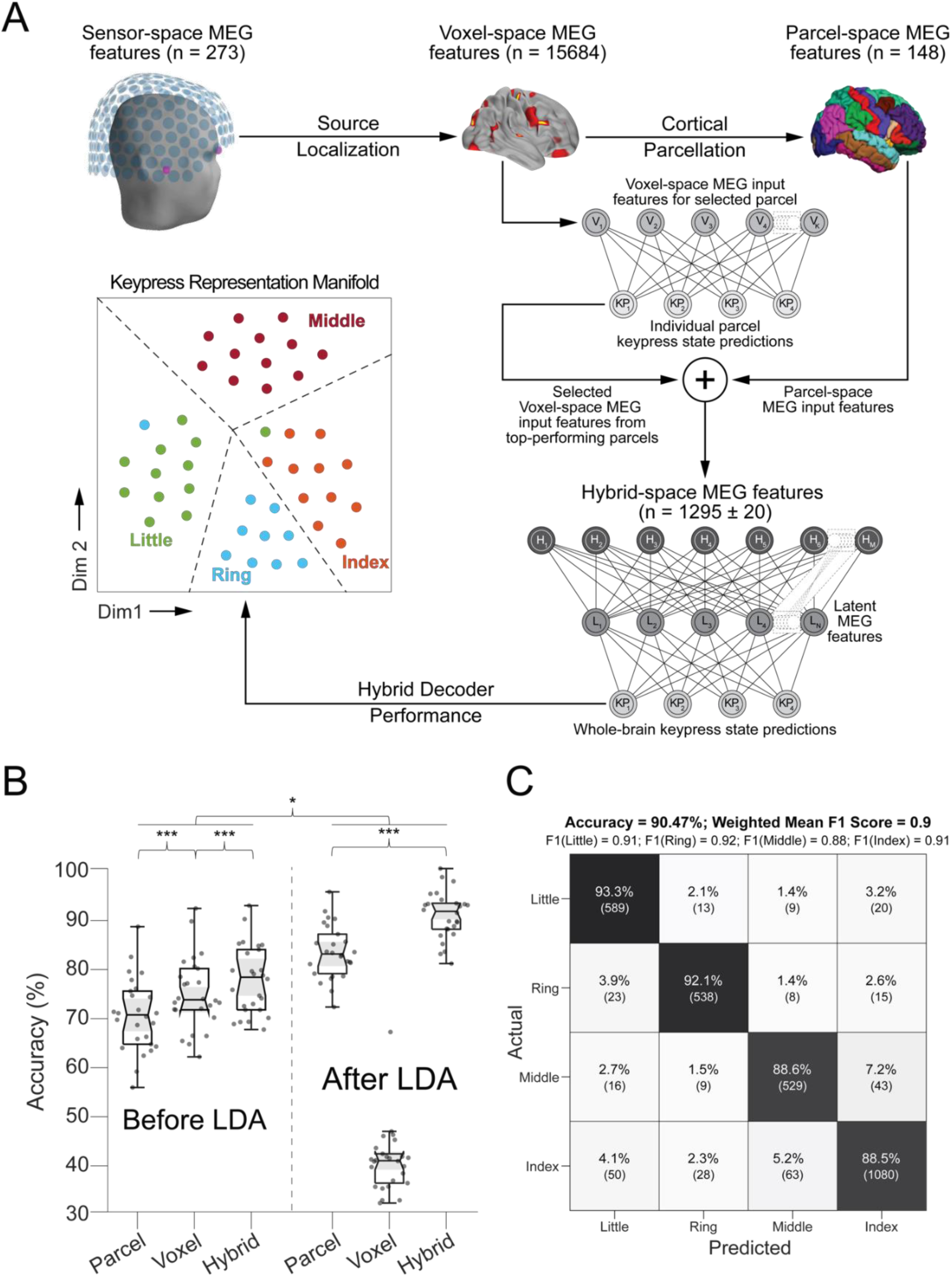
Hybrid spatial approach for neural decoding during skill learning *A. Pipeline*. Sensor-space MEG data (𝑁 = 272 channels) were source-localized (voxel-space features; 𝑁 = 15684 voxels), and then parcellated (parcel-space features; 𝑁 = 148) by averaging the activity of all voxels located within an individual region defined in a standard template space (Desikan-Killiany Atlas). Individual regional voxel-space decoders were then constructed and ranked. The final hybrid-space keypress state (i.e. – 4-class) decoder was constructed using all whole-brain parcel-space features and top-ranked regional voxel-space features (see Methods). **B. Decoding performance across parcel, voxel, and hybrid spaces.** Note that decoding performance was highest for the hybrid space approach compared to performance obtained for whole-brain voxel- and parcel spaces. Addition of linear discriminant analysis (LDA)-based dimensionality reduction further improved decoding performance for both parcel- and hybrid-space approaches. Each dot represents accuracy for a single participant and method. “**∗∗∗**” indicates 𝑝 < 0.001 and “**∗**” indicates 𝑝 < 0.05. ***C. Confusion matrix of individual finger identity decoding for hybrid-space manifold features.*** True predictions are located on the main diagonal. Off-diagonal elements in each row depict false-negative predictions for each finger, while off-diagonal elements in each column indicate false-positive predictions. Please note that the index finger keypress had the highest false-negative misclassification rate (11.55%).

We implemented different dimensionality reduction or manifold extraction strategies including principal component analysis (PCA), multi-dimensional scaling (MDS), minimum redundant maximum relevance (MRMR), and linear discriminant analysis (LDA) [34] to map the input feature (parcel, voxel, or hybrid) space to a low-dimensional latent space [18]. LDA-based manifold extraction led to the greatest classifier performance gains, improving keypress decoding accuracy to 90.47% ± SD 3.44% (**Figure 3B**; weighted mean F1 score = 0.91 ± SD 0.05). In comparison to the hybrid-space decoder, whole-brain parcel-space decoder performance also improved following LDA-based dimensionality reduction (82.95% ± SD 5.48%), while whole-brain voxel-space decoder accuracy dropped substantially (40.38 % ± SD 6.78%; also see **Figure 3 – figure supplement 2**).

Notably, decoding of index finger keypresses (executed at two different ordinal positions in the sequence) exhibited the highest false negative (0.115 per keypress) and false positive (0.067 per prediction) misclassification rates compared with all other digits (false negative rate range = [0.067 0.114]; false positive rate range = [0.085 0.131]; **Figure 3C**), raising the hypothesis that the same action could be differentially represented when executed within different contexts (i.e. – at different locations within the skill sequence). Testing the keypress state (4-class) hybrid decoder performance on Day 1 after randomly shuffling keypress labels for held-out test data resulted in a performance drop approaching expected chance levels (22.12%± SD 9.1%; **Figure 3 – figure supplement 3C**). An alternate decoder trained on ICA components labeled as movement or physiological artefacts (e.g. – head movement, ECG, eye movements and blinks; **Figure 3 – figure supplement 3A, D**) and removed from the original input feature set during the pre-processing stage approached chance-level performance (**Figure 4 – figure supplement 3)**, indicating that the 4-class hybrid decoder results were not driven by task-related artefacts.

Utilizing the highest performing decoders that included LDA-based manifold extraction, we assessed the robustness of hybrid-space decoding over multiple sessions by applying it to data collected on the following day during the *Day 2 Retest* (9-trial retest of the trained sequence) and *Day 2 Control* (single-trial performance of 9 different untrained sequences) blocks. The decoding accuracy for *Day 2* MEG data remained high (87.11% ± SD 8.54% for the trained sequence during *Retest*, and 79.44% ± SD 5.54% for the untrained *Control* sequences; **Figure 3 – figure supplement 4**). Thus, index finger classifiers constructed using the hybrid decoding approach robustly generalized from Day 1 to Day 2 across trained and untrained keypress sequences.

### Inclusion of keypress sequence context location optimized decoding performance

Next, we tracked the trial-by-trial evolution of keypress action manifolds as training progressed. Within-subject keypress neural representations progressively differentiated during early learning. A representative example in **Figure 4A** (top row) depicts increased four-digit representation clustering across trials 1, 11, and 36. The cortical representation of these clusters changed over the course of training, beginning with predominant involvement of contralateral pre-central areas in trial 1 before transitioning to greater contralateral post-central, superior frontal, and middle frontal cortex contributions in trials 11 and 36 (**Figure 4A**, bottom row), paralleling improvements in decoding performance (see **Figure 4 – figure supplement 1** for trial-by-trial quantitative feature importance score changes during skill learning).

**Figure 4.**
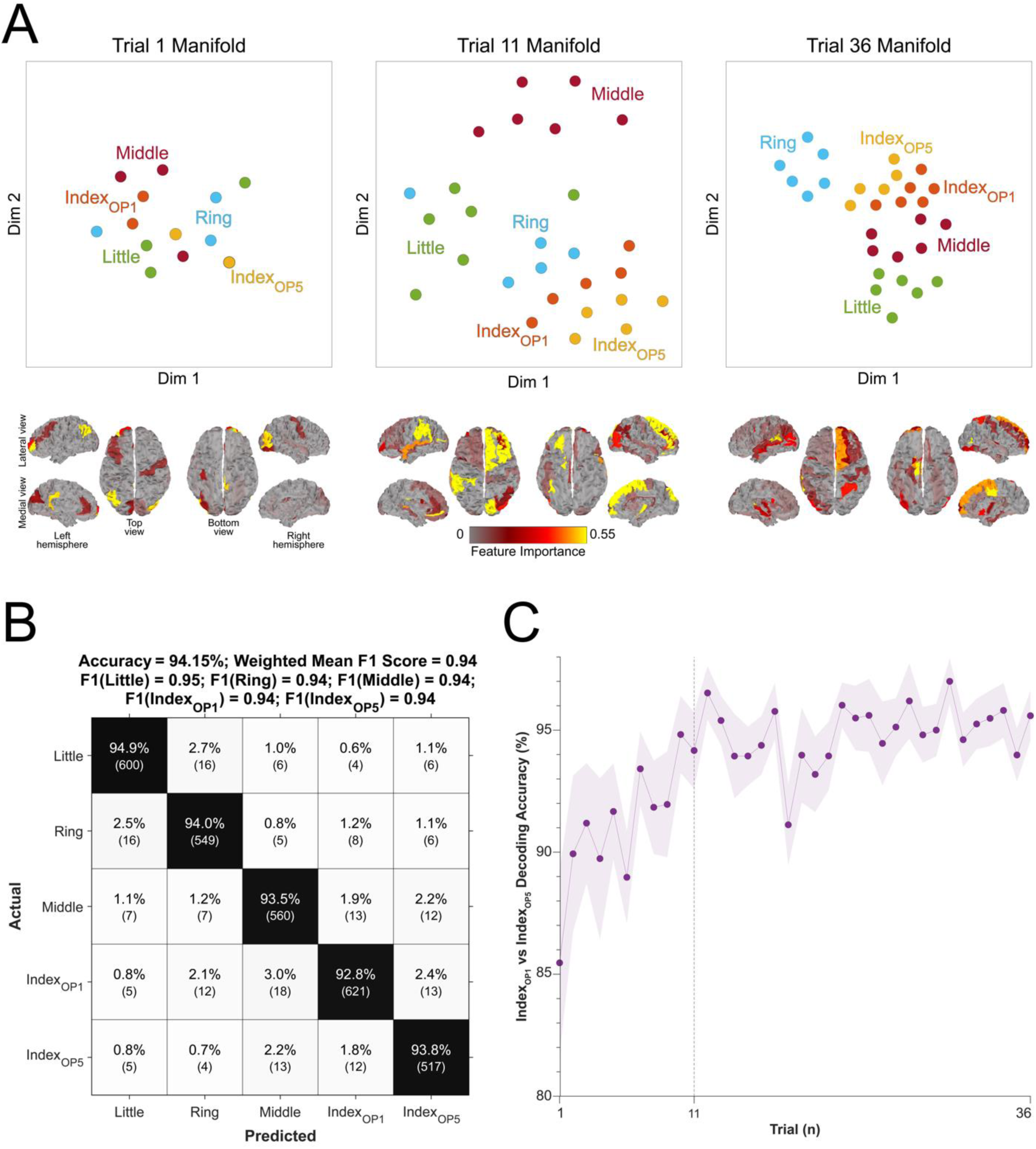
Evolution of keypress neural representations with skill learning. *A. Keypress neural representations differentiate during early learning.* t-SNE distribution of neural representation of each keypress (top scatter plots) is shown for trial 1 (start of training; top-left), 11 (end of early learning; top-center), and 36 (end of training; top-right) for a single representative participant. Individual keypress manifold representation clustering in trial 11 (top-center; end of early learning) depicts sub-clustering for the index finger keypress performed at the two different ordinal positions in the sequence (Index_OP1_ and Index_OP5_), which remains present by trial 36 (top-right). Spatial distribution of regional contributions to decoding (bottom brain surface maps). The surface color heatmap indicates feature importance scores across the brain. Note that decoding contributions shifted from contralateral right pre-central cortex at trial 1 (bottom-left) to contralateral superior and middle frontal cortex at trials 11 (bottom-center) and 36 (bottom-right). *B. Confusion matrix for 5-class decoding of individual sequence items.* Decoders were trained to classify contextual representations of the keypresses (i.e., 5-class classification of the sequence elements 4-1-2-3-4). Note that the decoding accuracy increased to 94.15% ± SD 4.84% and the misclassification of keypress 4 was significantly reduced (from 141 to 82). *C. Trial-by-trial classification accuracy for 2-class decoder (Index_OP1_ vs. Index_OP5_).* A decoder (200 ms window duration aligned to the KeyDown event) was trained to differentiate between the two index finger keypresses embedded at different positions within the practiced skill sequence (Index_OP1_ = index finger keypress at ordinal position 1 of the sequence; Index_OP5_ = index finger keypress at ordinal position 5 of the sequence). Decoder accuracy progressively improved over early learning, stabilizing around 96% by trial 11 (end of early learning). Similar results were observed for other decoding window sizes (50, 100, 150, 250 and 300 ms; see **Figure 4 – figure supplement 2**). Taken together, these findings indicate that the neural feature space evolves over early learning to incorporate sequence location information.

The trained skill sequence required pressing the index finger twice (**4**-1-3-2-**4**) at two contextually different ordinal positions (sequence positions 1 and 5). Inclusion of sequence location information (i.e. – sequence context) for each keypress action (five sequence elements with the one keypress represented twice at two different locations) improved decoding accuracy (*t* = 7.09, *p* < 0.001, *df* = 25, Cohen’s *d* = 0.86, **Figure 4B)** from 90.47% (± SD 3.44%) to 94.15% (± SD 4.84%; weighted mean F1 score: 0.94), and reduced overall misclassifications by 54.3% (from 219 to 119; **Figure 3C and 4B**). The improved decoding accuracy is supported by greater differentiation in neural representations of the index finger keypresses performed at positions 1 and 5 of the sequence (**Figure 4A**), and by the trial-by-trial increase in 2-class decoding accuracy over early learning (**Figure 4C)** across different decoder window durations (**Figure 4 – figure supplement 2**). As expected, the 5-class hybrid-space decoder performance approached chance levels when tested with randomly shuffled keypress labels (18.41%± SD 7.4% for Day 1 data; **Figure 4 – figure supplement 3C**). Task-related eye movements did not explain these results since an alternate 5-class decoder constructed from three eye movement features (gaze position at the KeyDown event, gaze position 200ms later, and peak eye movement velocity within this window; **Figure 4 – figure supplement 3A**) performed at chance levels (cross-validated test accuracy = 0.2181; **Figure 4 – figure supplement 3B, C)**.

On Day 2, incorporating contextual information into the hybrid-space decoder enhanced classification accuracy for the trained sequence only (improving from 87.11% for 4-class to 90.22% for 5-class), while performing at or below-chance levels for the control sequences (≤ 30.22% ± SD 0.44%). Thus, the accuracy improvements resulting from inclusion of contextual information in the decoding framework was specific for the trained skill sequence.

### Neural representation of keypress sequence location diverged during early skill learning

We used a Euclidian distance measure to evaluate the differentiation of the neural representation manifold of the same action (i.e. - an index-finger keypress) executed within different local sequence contexts (i.e. - ordinal position 1 vs. ordinal position 5; **Figure 5**). To make these distance measures comparable across participants, a new set of classifiers was then trained with group-optimal parameters (i.e. – broadband hybrid-space MEG data with subsequent manifold extraction (**Figure 3 – figure supplements 2)** and LDA classifiers (**Figure 3 – figure supplements 7)** trained on 200ms duration windows aligned to the KeyDown event (see Methods, **Figure 3 – figure supplements 5**).

**Figure 5:**
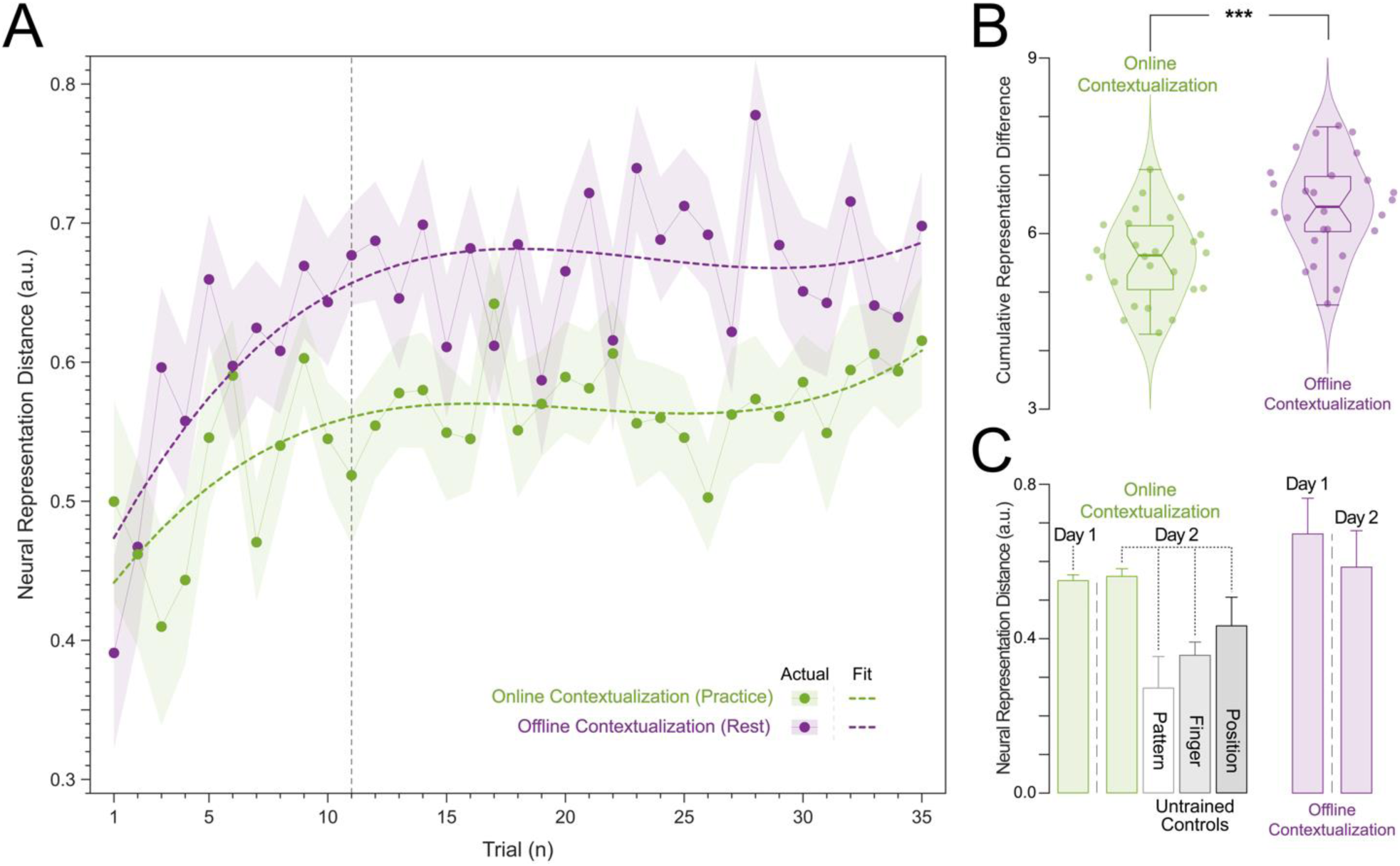
Neural representation distance between index finger keypresses performed at two different ordinal positions within a sequence. A. *Contextualization increases over Early Learning during Day 1 Training.* Online (green) and offline (purple) neural representation distances (*contextualization*) between two index finger key presses performed at ordinal positions 1 and 5 of the trained sequence (4-1-3-2-4) are shown for each trial during Day 1 Training. Both online and offline contextualization between the two index finger representations increase sharply over *Early Learning* before stabilizing across later *Day 1 Training* trials. B. *Contextualization develops predominantly during rest periods (offline) on Day 1.* The cumulative neural representation differences during early learning were significantly greater over rest (*Offline contextualization*; right) than during practice (*Online contextualization*; left) periods (*t* = 4.84, *p* < 0.001, *df* = 25, Cohen’s *d*= 1.2). C. *Contextualization acquired on Day 1 was retained on Day 2 specifically for the trained sequence.* The neural representation differences assessed across both rest and practice for the trained sequence (4-1-3-2-4) were retained at *Day 2 Retest*. This is in stark contrast with the reduction in contextualization for several untrained sequences controlling for: 1) index finger keypresses located at the same ordinal positions 1 and 5 but within a different intervening sequence pattern (*Pattern Specificity Control*: 4-2-3-1-4, 51.05% lower contextualization); 2) use of a finger different than the index (little or ring finger) in both ordinal positions 1 and 5 (*Finger Specificity Control*: 2-1-3-4-2, 1-4-2-3-1 and 2-3-1-4-2; 35.80% lower contextualization); and 3) multiple index finger keypresses occurring at ordinal positions other than 1 and 5 (*Position Specificity Control*: 4-2-4-3-1 and 1-4-3-4-2; 22.06% lower contextualization). Note that offline contextualization cannot be measured for the *Day 2 Control* sequences as each sequence was only performed over a single trial.

The Euclidian distance between neural representations of Index_OP1_ (i.e. - index finger keypress at ordinal position 1 of the sequence) and Index_OP5_ (i.e. - index finger keypress at ordinal position 5 of the sequence) increased progressively during early learning (**Figure 5A**)—predominantly during rest intervals (*offline contextualization*) rather than during practice (*online*) (*t* = 4.84, *p* < 0.001, *df* = 25, Cohen’s *d* = 1.2; **Figure 5B**; **Figure 5 – figure supplement 1A**). An alternative online contextualization determination equaling the time interval between online and offline comparisons (*Trial-based;* 10 seconds between Index_OP1_ and Index_OP5_ observations in both cases) rendered a similar result (**Figure 5 – figure supplement 2B**).

Offline contextualization strongly correlated with cumulative micro-offline gains (*r* = 0.903, *R²* = 0.816, *p* < 0.001; **Figure 5 – figure supplement 1A, inset**) across decoder window durations ranging from 50 to 250ms (**Figure 5 – figure supplement 1B, C)**. The offline contextualization between the final sequence of each trial and the second sequence of the subsequent trial (excluding the first sequence) yielded comparable results. This indicates that pre-planning at the start of each practice trial did not directly influence the offline contextualization measure [30] (**Figure 5 – figure supplement 2A**, *1^st^ vs. 2^nd^ Sequence approaches*). Conversely, online contextualization (using either measurement approach) did not explain early online learning gains (i.e. – **Figure 5 – figure supplement 3)**. Within-subject correlations were consistent with these group-level findings. The average correlation between offline contextualization and micro-offline gains within individuals was significantly greater than zero (**Figure 5 – figure supplement 4, left**; *t* = 3.87, *p* = 0.00035, *df* = 25, Cohen’s *d* = 0.76) and stronger than correlations between online contextualization and either micro-online (**Figure 5 – figure supplement 4, middle**; *t* = 3.28, *p* = 0.0015, *df* = 25, Cohen’s *d* = 1.2) or micro-offline gains (**Figure 5 – figure supplement 4, right**; *t* = 3.7021, *p* = 5.3013e-04, *df* = 25, Cohen’s *d* = 0.69). These findings were not explained by behavioral changes of typing rhythm (*t* = -0.03, *p* = 0.976; **Figure 5 – figure supplement 5**), adjacent keypress transition times (*R^2^* = 0.00507, F [1,3202] = 16.3; **Figure 5 – figure supplement 6**), or overall typing speed (between-subject; *R^2^* = 0.028, *p* = 0.41; **Figure 5 – figure supplement 7**).

Finally, contextualization of Index_OP1_ vs. Index_OP5_ representations observed on *Day 1* generalized to *Day 2 Retest* of the trained skill sequence. Distances between representations for the same keypress performed twice within untrained sequences were lower in magnitude (*Day 2 Control*)— pointing to specificity of the contextualization effect **(Figure 5C).**

## Discussion

The main findings of this study during which subjects engaged in a naturalistic, self-paced task were that individual sequence action representations differentiate during early skill learning in a manner reflecting the local sequence context in which they were performed, and that the degree of representational differentiation—particularly prominent over rest intervals—correlated with skill gains.

### Optimizing decoding of sequential finger movements from MEG activity

The initial phase of the study focused on optimizing the accuracy of decoding individual finger keypresses from MEG brain activity. Recent work showed that the brain simultaneously processes information more efficiently across multiple—rather than a single—spatial scale(s) [28, 32]. To this effect, we developed a novel hybrid-space approach designed to integrate neural representation dynamics over two different spatial scales: (1) *whole-brain parcel-space* (i.e. – spatial activity patterns across all cortical brain regions) and (2) *regional voxel-space* (i.e. – spatial activity patterns within select brain regions) activity. We found consistent spatial differences between whole-brain parcel-space feature importance (predominantly contralateral frontoparietal, **Figure 2B**) and regional voxel-space decoder accuracy (bilateral sensorimotor regions, **Figure 2D**). The *whole-brain parcel-space* decoder likely emphasized more stable activity patterns in contralateral frontoparietal regions that differed between individual finger movements [21, 35], while the *regional voxel-space* decoder likely incorporated information related to adaptive interhemispheric interactions operating during motor sequence learning [32, 36, 37], particularly pertinent when the skill is performed with the non-dominant hand [38–40]. The observation of increased cross-validated test accuracy (as shown in **Figure 3 – Figure Supplement 6**) indicates that the spatially overlapping information in parcel- and voxel-space time-series in the hybrid decoder was complementary, rather than redundant [41]. The hybrid-space decoder which achieved an accuracy exceeding 90%—and robustly generalized to Day 2 across trained and untrained sequences—surpassed the performance of both parcel-space and voxel-space decoders and compared favorably to other neuroimaging-based finger movement decoding strategies [6, 24, 42–44].

Evaluation of individual brain oscillatory activity revealed that low-frequency oscillations (LFOs) result in higher decoding accuracy compared to other narrow-band activity [18, 45]. Task-related movements—which also express in lower frequency ranges—did not explain these results given the near chance-level performance of alternative decoders trained on (a) artefact-related ICA components removed during MEG pre-processing (**Figure 3 – figure supplement 3A-C)** and on (b) task-related eye movement features **(Figure 4 – figure supplement 3B, C)**. This explanation is also inconsistent with the minimal average head motion of 1.159 mm (± 1.077 SD) across the MEG recording (**Figure 3 – figure supplement 3D)**. How could LFOs contribute to keypress decoding accuracy? LFOs, observed during movement onset in the cerebral cortex of animals [46, 47] and humans [48–50], encode information about movement trajectories and velocity [46, 47]. They also contain information related to movement timing [51–53], preparation [54, 55], sensorimotor integration [49], kinematics [54, 55] and may contribute to the precise temporal coordination of movements required for sequencing [56]. Within clinical contexts, LFOs in the frontoparietal regions, resulting in high decoding accuracy in the present study, have been linked to recovery of motor function after brain lesions like stroke [48, 51, 57].

### Neural representations of individual sequence actions differentiate during early skill learning

Next, we exploited the hybrid decoding approach to investigate if individual sequence action representations differentiate or remain stable during early skill learning, when the memory is not yet fully formed [1]. The first hint of representational differentiation was the highest false-negative and lowest false-positive misclassification rates for index finger keypresses performed at different locations in the sequence compared with all other digits (**Figure 3C**). This was further supported by the progressive differentiation of neural representations of the index finger keypress (**Figure 4A**) and by the robust trial-by-trial increase in 2-class (Index_OP1_ vs Index_OP5_) decoding accuracy across time windows ranging between 50 and 250ms (**Figure 4C; Figure 4 – figure supplement 2**). Further, the 5-class classifier—which directly incorporated information about the sequence location context of each keypress into the decoding pipeline—improved decoding accuracy relative to the 4-class classifier (**Figure 4C**). Importantly, testing on Day 2 revealed specificity of this representational differentiation for the trained skill but not for the same keypresses performed during various unpracticed control sequences (**Figure 5C**).

The main region contributing information to representational differentiation during early practice (trials 1-10) was the primary motor cortex, followed by the somatosensory cortex (trial 11), both of which are known to be actively engaged in skill acquisition [6, 16, 58–61]. Concurrently, information from the superior frontal and middle frontal cortex—which encode hierarchical structures of skill sequences [20]—steadily increased in importance and emerged as the two most crucial decoding contributors once skill performance plateau had been reached (trials 15 to 36; **Figure 4 – figure supplement 1)** [27, 62]. Thus, the neural substrates supporting finger movements and their representational differentiation during early skill learning (the time period during which 95% skill gains in the training session occur [1, 29], trials 1-11 in this study) differed from those supporting stable performance during the subsequent skill plateau period [16, 63] (trials 12-36 in this study).

### Differentiation of neural representations developed predominantly during rest periods interspersed with practice

We then focused on the timeline of differentiation of index finger keypress neural representations—which we refer to as contextualization—over early learning. We found that contextualization increased progressively during early learning—predominantly during short rest breaks (offline) rather than during practice (online; **Figure 5** and **Figure 5 – figure supplement 2B**). Offline contextualization consistently correlated with early learning gains across a range of decoding windows (50–250ms; **Figure 5 – figure supplement 1**). This result remained unchanged when measuring offline contextualization between the last and second sequence of consecutive trials, inconsistent with a possible confounding effect of pre-planning [30] (**Figure 5 – figure supplement 2A**). On the other hand, online contextualization did not predict learning (**Figure 5 – figure supplement 3**). Consistent with these results the average within-subject correlation between offline contextualization and micro-offline gains was significantly stronger than within-subject correlations between online contextualization and either micro-online or micro-offline gains (**Figure 5 – figure supplement 4)**.

Offline contextualization was not driven by trial-by-trial behavioral differences, including typing rhythm (**Figure 5 – figure supplement 5**) and adjacent keypress transition times (**Figure 5 – figure supplement 6**) nor by between-subject differences in overall typing speed (**Figure 5 – figure supplement 7**)—ruling out a reliance on differences in the temporal overlap of keypresses. Importantly, offline contextualization documented on Day 1 stabilized once a performance plateau was reached (trials 11-36), and was retained on Day 2, documenting overnight consolidation of the differentiated neural representations. A possible neural mechanism supporting contextualization could be the emergence and stabilization of conjunctive “what–where” representations of procedural memories [64] with the corresponding modulation of neuronal population dynamics [65, 66] during early learning. Exploring the link between contextualization and neural replay could provide additional insights into this issue [6, 12, 13, 15].

In this study, classifiers were trained on MEG activity recorded during or immediately after each keypress, emphasizing neural representations related to action execution, memory consolidation and recall over those related to planning. An important direction for future research is determining whether separate decoders can be developed to distinguish the representations or networks separately supporting these processes. Ongoing work in our lab is addressing this question. The present accuracy results across varied decoding window durations and alignment with each keypress action support the feasibility of this approach (**Figure 3—figure supplement 5**).

## Limitations

One limitation of this study is that contextualization was investigated for only one finger movement (index finger or digit 4) embedded within a relatively short 5-item skill sequence. Determining if representational contextualization is exhibited across multiple finger movements embedded within for example longer sequences (e.g. – two index finger and two little finger keypresses performed within a short piece of piano music) will be an important extension to the present results. While a supervised manifold learning approach (LDA) was used here because it optimized hybrid-space decoder performance, unsupervised strategies (e.g. - PCA and MDS, which also substantially improved decoding accuracy in the present study; **Figure 3 – figure supplement 2**) are likely more suitable for real-time BCI applications. Finally, caution should be exercised when extrapolating findings during early skill learning, a period of steep performance improvements, to findings reported after insufficient practice [67], post-plateau performance periods [68], or non-learning situations (e.g. performance of non-repeating keypress sequences in [67]) when reactive inhibition or contextual interference effects are prominent. Ultimately, it will be important to develop new paradigms allowing one to independently estimate the different coincident or antagonistic features (e.g. - memory consolidation, planning, working memory and reactive inhibition) contributing to micro-online and micro-offline gains during and after early skill learning within a unifying framework.

## Summary

In summary, individual sequence action representations contextualize during early learning of a new skill and the degree of differentiation parallels skill gains. Differentiation of the neural representations developed during rest intervals of early learning to a larger extent than during practice in parallel with rapid consolidation of skill. It is possible that the systematic inclusion of contextualized information into sequence skill practice environments could improve learning in areas as diverse as music education, sports training, and rehabilitation of motor skills after brain lesions.

## Materials and Methods

### Study Participants

The study was approved by the Combined Neuroscience Institutional Review Board of the National Institutes of Health (NIH). A total of thirty-three young and healthy adults (16 females) with a mean age of 26.6 years (± 0.87 SEM) participated in the study after providing written informed consent and undergoing a standard neurological examination. No participants were actively engaged in playing musical instruments in their daily lives, as per guidelines outlined in prior research [69, 70]. All study scientific data were de-identified and permanently unlinked from all personal identifiable information (PII) before the analysis. These data are publicly available upon request (https://nih.box.com/v/hcpsSkillLearningData). Two participants were excluded from the analysis due to MEG system malfunction during data acquisition. An additional 5 subjects were excluded because they failed to generate any correct sequences in two or more consecutive trials. The study was powered to determine the minimum sample size needed to detect a significant change in skill performance following training using a one-sample t-test (two-sided; alpha = 0.05; 95% statistical power; Cohen’s *d* effect size = 0.8115 calculated from previously acquired data in our lab [71]). The calculated minimum sample size was 22. The included study sample size (n = 26) exceeded this minimum [1].

### Experimental Setup

Participants practiced a procedural motor skill learning task that involved repetitively typing a 5-item numerical sequence (4-1-3-2-4) displayed on a computer screen. They were instructed to perform the task as quickly and accurately as possible on a response pad (Cedrus LS-LINE, Cedrus Corp) using their non-dominant, left hand. Each numbered sequence item corresponded to a specific finger keypress: 1 for the little finger, 2 for the ring finger, 3 for the middle finger, and 4 for the index finger. Individual keypress times and identities were recorded and used to assess skill learning and performance. The response pad was positioned in a manner that minimized wrist, arm or more proximal body movements during the task. The head was restrained with an inflatable air bladder, and head position was assessed at the beginning and at the end of each recording. The mean measured head movement across the study group was 1.159 mm (± 1.077 SD; **Figure 3 – figure supplement 3**)

Participants practiced the skill for 36 trials. Each trial spanned a total of 20 seconds and included a 10-second practice round followed by a 10-second rest break. The study design followed specific recommendations by Pan and Rickard (2015): 1) utilizing 10-second practice trials and 2) constraining analysis of micro-offline gains to early learning trials (where performance monotonically increases and 95% of overall performance gains occur) that precede the emergence of “scalloped” performance dynamics strongly linked to reactive inhibition effects ([29, 72]). This is precisely the portion of the learning curve Pan and Rickard referred to when they stated “…*rapid learning during that period masks any reactive inhibition effect”* [29].

The five-item sequence was displayed on the computer screen for the duration of each practice round and participants were directed to fix their gaze on the sequence. Small asterisks were displayed above a sequence item after each successive keypress, signaling the participants’ present position within the sequence. Inclusion of this feedback minimizes working memory loads during task performance [73]. Following the completion of a full sequence iteration, the asterisk returned to the first sequence item. The asterisk did not provide error feedback as it appeared for both correct and incorrect keypresses. At the end of each practice round, the displayed number sequence was replaced by a string of five “X” symbols displayed on the computer screen, which remained for the duration of the rest break. Participants were instructed to focus their gaze on the screen during this time. The behavior in this explicit, motor learning task consists of generative action sequences rather than sequences of stimulus-induced responses as in the serial reaction time task (SRTT). A similar real-world example would be manually inputting a long password into a secure online application in which one intrinsically generates the sequence from memory and receives similar feedback about the password sequence position (also provided as asterisks), which is typically ignored by the user.

On the next day, participants were tested (*Day 2 Retest*) with the same trained sequence (4-1-3-2-4) for 9 trials as well as for 9 different unpracticed control sequences (*Day 2 Control*; 2-1-3-4-2; 4-2-4-3-1; 3-4-2-3-1; 1-4-3-4-2; 3-2-4-3-1; 1-4-2-3-1; 3-2-4-2-1; 2-3-1-4-2; 4-2-3-1-4) each for one trial. The practice schedule structure for Day 2 was same as Day 1, with 10-second practice trials interleaved with 10-second rest breaks.

### Behavioral data analysis

#### Skill

Skill, in the context of the present task, is quantified as the *correct sequence typing speed*, (i.e., - the number of correctly typed sequence keypresses per second; kp/s). That is, improvements in the speed/accuracy trade-off equate to greater skill. Keypress transition times (KTT) were calculated as the difference in time between the *keyDown* events recorded for consecutive keypresses. Since the sequence was repeatedly typed within a single trial, individual keypresses were marked as correct if they were members of a 5 consecutive keypress set that matched any possible circular shift of the displayed 5-item sequence. The instantaneous correct sequence speed was calculated as the inverse of the average KTT across a single correct sequence iteration and was updated for each correct keypress. Trial-by-trial skill changes were assessed by computing the median correct sequence typing speed for each trial.

#### Early Learning

The *early learning* period was defined as the trial range (1 - T trials) over which 95% of the total skill performance was first attained at the group level. We quantified this by fitting the group average trial-by-trial correct sequence speed data with an exponential model of the form:

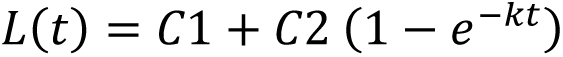

Here, the trial number is denoted by *t*, and *L(t)* signifies the group-averaged performance at trial *t*. Parameters *C1* and *C2* correspond to the pre-training performance baseline and asymptote, respectively, while *k* denotes the learning rate. The values for *C1*, *C2*, and *k* were computed using a constrained nonlinear least-squares method (MATLAB’s lsqcurvefit function, trust-region-reflective algorithm) and were determined to be 0.5, 0.15, and 0.2, respectively. The early learning trial cut-off, denoted as *T*, was identified as the first trial where 95% of the learning had been achieved. In this study, *T* was determined to be trial 11.

#### Micro-offline and -online gains

Performance improvements over each 10-second rest break (*micro-offline gains)* were calculated as the net performance change (instantaneous correct sequence typing speed) from the end of one practice period to the onset of the next, while *micro-online gains* were computed as the net performance change over a single practice trial. Total early learning was derived as the sum of all *micro-online* and *micro-offline* gains over trials 1-11. Cumulative micro-offline gains, micro-online gains, and total early learning were statistically compared using 1-way ANOVAs and post-hoc Tukey tests. Possible pre-planning effects on initial skill performance [30] were assessed by using paired t-tests to statistically compare cumulative micro-offline and -online computed for all keypresses with their measurement counterparts calculated after omitting the first 3 keypresses in each trial from the correct sequence speed computation.

### MRI Acquisition

We acquired T1-weighted high-resolution anatomical MRI volumes images (1 mm^3^ isotropic MPRAGE sequence) for each participant on a 3T MRI scanner (GE Excite HDxt or Siemens Skyra) equipped with a 32-channel head coil. These data allowed for spatial co-registration of an individual participant’s brain with the MEG sensors, and individual head models required for surface-based cortical dipole estimation from MEG signals (i.e. – MEG source-space modeling).

### MEG Acquisition

We recorded continuous magnetoencephalography (MEG) at a sampling frequency of 600 Hz using a CTF 275 MEG system (CTF Systems, Inc., Canada) while participants were seated in an upright position. The MEG system comprises a whole-head array featuring 275 radial 1^st^-order gradiometer/SQUID channels housed in a magnetically shielded room (Vacuumschmelze, Germany). Three of the gradiometers (two non-functional and one with high channel noise after visual inspection) were excluded from the analysis resulting in a total of 272 useable MEG sensor channels. Synthetic 3^rd^ order gradient balancing was applied to eliminate background noise in real-time data collection. Temporal alignment of behavioral and MEG data was achieved using a TTL trigger. Head position in the scanner coordinate space was assessed at the beginning and end of each recording using head localization coils at the nasion, left, and right pre-auricular locations. These fiducial positions were co-registered in the participants’ T1-MRI coordinate space using a stereotactic neuronavigation system (BrainSight, Rogue Research Inc.). MEG data was acquired starting 6 min before the task (resting-state baseline) and continued through the end of the 12 min training session.

### MEG Data Analysis

#### Preprocessing

MEG data were preprocessed using the FieldTrip [74] and EEGLAB [75] toolboxes on MATLAB 2022a. Continuous raw MEG data were band-pass filtered between 1-100 Hz with a 4^th^ order noncausal Butterworth filter. 60 Hz line noise was removed with a narrow-band discrete Fourier transform (DFT) notch filter. Independent component analysis (ICA) was used to remove typical MEG signal artifacts associated with eye blinks or movement, muscle contraction or cardiac pulsation. All recordings were visually inspected and marked to denoise segments containing other large amplitude artifacts due to movements. Eye movements were simultaneously recorded with MEG (EyeLink 1000 Plus).

#### Source Reconstruction and Parcellation

For each participant, individual volume conduction models were computed to estimate the propagation of brain-generated currents through tissue resulting in externally measurable magnetic fields. This was accomplished through a single-shell head corrected-sphere approach based on the brain volume segmentation of the participant’s high-resolution T1 MRI. Source models and surface labels from the Desikan-Killiany Atlas [31] were created for each participant using inner-skull and pial layer surfaces obtained through FreeSurfer segmentation [76, 77] and Connectome Workbench resampling [76, 77]. Aligning sensor positions in the MEG helmet to individual head space involved rigid-body registration of the mean MEG head coil position to the same fiducial locations marked in the MRI and applying the same affine transformation to all MEG sensors.

The individual source, volume conduction model, and sensor positions were then utilized to generate the forward solution at each source dipole location, describing the propagation of source activity from each cortical location on the grid to each MEG sensor. The Linearly Constrained Minimum-Variance (LCMV) beamformer was employed for computing the inverse solution. Each trial of MEG activity contributed to calculating the inverse solution data covariance matrix. The individual sample noise covariance matrix was derived from 6 minutes of pre-training rest MEG data recorded in the same subject during the same session. A total of 15,684 surface-based cortical dipoles (i.e. - source-space voxels) were estimated.

Source-space parcellation was carried out by averaging all voxel time-series located within distinct anatomical regions defined in the Desikan-Killiany Atlas [31]. Since source time-series estimated with beamforming approaches are inherently sign-ambiguous, a custom Matlab-based implementation of the *mne.extract_label_time_course* with “*mean_flip”* sign-flipping procedure in MNE-Python [78] was applied prior to averaging to prevent within-parcel signal cancellation. All voxel time-series within each parcel were extracted and the time-series sign was flipped at locations where the orientation difference was greater than 90° from the parcel mode. A mean time-series was then computed across all voxels within the parcel after sign-flipping.

### Feature Selection for Decoding

Several MEG activity features were extracted over different spatial, spectral, and temporal scales.

#### Oscillatory Analysis

MEG signals were constrained to broadband (1-100 Hz) or standard neural oscillatory frequency bands defined as delta (1-3 Hz), theta (4-7 Hz), alpha (8-14 Hz), beta (15-24 Hz), gamma (25-50 Hz), and high-gamma (51-100 Hz) using a 4^th^ order non-causal Butterworth filter. Subsequent decoding analyses were independently conducted for each band of MEG activity.

#### Spatial Analysis

Decoding was performed in both sensor and source spaces. The sensor-space decoding feature dimension was 272 (corresponding to the 272 useable MEG channels), while source-space decoding was carried out at both the higher feature dimension voxel (i.e. – higher spatially sampled; N = 15,684) and lower feature dimension parcel space (i.e. – lower spatially sampled; N = 148) across all oscillatory frequency bands (i.e. - broadband, delta, theta, alpha, beta, gamma, and high-gamma) for comprehensive comparison.

#### Temporal Analysis

MEG activity time-series corresponding to each keypress was defined using the time window, [t + Δt], where t ɛ [0 : 10 ms : 100 ms] and Δt ɛ [25 ms : 25 ms : 350 ms]. In other words, a sliding window of variable width (from 25 ms to 350 ms with 25 ms increments), and with onsets ranging from the *keyDown* event (i.e. – t = 0 ms) to +100 ms after the *keyDown* event (with increments of 10 ms) was used. This approach generated a set of 140 different time windows associated with each keypress for each participant. MEG activity was averaged over time within each of these widows and independently analyzed for decoding. The optimal time window was selected for each subject that resulted in the maximum cross-validation performance (**Figure 3 – figure supplement 5**). This window optimization analysis was performed for each frequency band and spatial scale.

#### Hybrid Spatial Approach

First, we evaluated the decoding performance of each individual brain region in accurately decoding finger keypresses from regional voxel-space (i.e. - all voxels within a brain region as defined by the Desikan-Killiany Atlas) activity. Brain regions were then ranked from 1 to 148 based on their decoding accuracy at the group level. In a stepwise manner, we then constructed a “hybrid-space” decoder by incrementally concatenating regional voxel-space activity of brain regions—starting with the top-ranked region—with whole-brain parcel-level features and assessed decoding accuracy. Subsequently, we added the regional voxel-space features of the second-ranked brain region and continued this process until decoding accuracy reached saturation. The optimal “hybrid-space” input feature set over the group included the 148 parcel-space features and regional voxel-space features from a total of 8 brain regions (bilateral superior frontal, middle frontal, pre-central and post-central; N = 1295 ± 20 features).

#### Dimension Reduction

We independently applied several supervised and unsupervised dimension reduction techniques as an additional feature extraction step for each broadband MEG activity space (i.e. - sensor, parcel, voxel, and hybrid), including: linear discriminant analysis (LDA), minimum redundant maximum relevance (MRMR), principal component analysis (PCA), Autoencoder, Diffusion maps, factor analysis, large margin nearest neighbor (LMNN), multi-dimensional scaling (MDS), neighbor component analysis (NCA), spatial predictor envelope (SPE)[34]. Among these techniques, PCA, MDS, MRMR, and LDA emerged as particularly effective in significantly improving decoding performance across all broadband MEG activity spaces.

PCA, a method for unsupervised dimensionality reduction, transforms the high-dimensional dataset into a new coordinate system of orthogonal principal components. These components, capturing the maximum variance in the data, were iteratively added to reconstruct the feature space and execution of decoding. MDS finds a configuration of points in a lower-dimensional space such that the distances between these points reflect the dissimilarities or similarities between the corresponding objects in the original high-dimensional space. MRMR, an approach combining relevance and redundancy metrics, ranks features based on their significance to the target variable and their non-redundancy with other features. The decoding process started with the highest-ranked feature and iteratively incorporated subsequent features until decoding accuracy reached saturation. LDA finds the linear combinations of features (dimensions) that best separate different classes in a dataset. It projects the original features onto a lower-dimensional space (number of classes -1) while preserving the class-discriminatory information. This transformation maximizes the ratio of the between-class variance to the within-class variance. In our study, LDA transformed the features to a 3/4-dimensional hyperdimensional space that were used for decoding. Dimension reduction was first applied to training data, and then with the tuned parameters of the dimension reduction model, an independent test data subset was transformed for decoder metrics evaluation. Decoding accuracies were systematically compared between the original and reduced dimension feature spaces, providing insight into the effectiveness of each dimension reduction technique. By rigorously assessing the impact of dimension reduction on decoding accuracy, the study aimed to identify techniques that not only reduced the computational burden associated with high-dimensional data but also enhanced the discriminative power of the selected features. This comprehensive approach aimed at optimizing the neural representation of each finger keypress for decoding performance across various spatial contexts.

### Decoding Analysis

Decoding analysis was conducted for each participant individually, employing a randomized split of the data into independent training (90%) and test (10%) samples over eight iterations. For each iteration, an 8-fold cross-validation was applied to the training samples to optimize decoder configuration, allowing for the fine-tuning of hyperparameters and selection of the most effective model. On average, the total number of individual keypress samples for the entire duration of training was 219 ± SD: 66 (keypress 1: little), 205 ± SD: 66 (keypress 2: ring), 209 ± 66 (keypress 3: middle), and 426 ± SD: 131 (keypress 4: index) across participants. Only keypresses belonging to correctly typed sequence iterations (94.64% ± 4.04% of all keypresses) were considered. The total number of index finger keypresses (i.e. - keypress 4) was approximately twice that of the others, as it was the only action that occurred more than once in the trained sequence (4-1-3-2-4), albeit at two different ordinal positions. Considering the higher (2x) number of samples for one-class, we independently oversampled the keypresses 1, 2 and 3 to avoid overfitting to the over-represented class. Importantly, oversampling was applied independently for each keypress class, ensuring that validation folds were never oversampled, and training folds did not share common oversampled patterns. The decoder configuration demonstrating the best validation performance was selected for each iteration, and subsequently, test accuracy was evaluated on the independent/unseen test samples. This process was repeated for the 8 different iterations of train-test splitting, and the average test accuracy over all iterations was reported. This rigorous methodology aimed at generalizing decoding performance to ensure robust and reliable results across participants. Finally, decoding evaluation was also performed on the Day 2 data, for both the trained (*Day 2 Retest;* 9 trials) and untrained sequences (*Day 2 Control*; 9 different single-trial tests).

#### Machine Learning Classifiers

We employed a diverse set of machine learning-based decoders—including Naïve Bayes (NB), decision trees (DT), ensembles (EN), k-nearest neighbor (KNN), linear discriminant analysis (LDA), support vector machines (SVM), and artificial neural network (ANN)—to train features generated over all possible combinations of spatial and temporal scales and oscillation frequency-bands in order to carry out a comprehensive comparative analysis. The hyperparameters of these decoders underwent fine-tuning using Bayesian optimization search.

All NB classifiers were configured with a normal distribution predictor and Gaussian Kernel, while KNN classifiers had a K value of 4 (for keypress decoding) and utilized the Euclidean distance metric. For DT classifiers, the maximum number of splits was set to 4 (for keypress decoding), with leaves being merged based on the sum of risk values greater or equal to the risk associated with the parent node. The optimal sequence of pruned trees was estimated, and the predictor selection method was ’Standard CART,’ selecting the split predictor that maximizes the split-criterion gain over all possible splits of all predictors. The split criterion used was ’gdi’ (Gini’s diversity index). EN classifiers employed the bagging method with random predictor selections at each split, forming a random forest. The maximum number of learning cycles was set to 100 with a weak learner based on discriminant analysis. For SVM, the RBF kernel was selected through cross-validation (CV), and the ’C’ parameter and kernel scale were optimized using Bayesian optimization search. In the case of LDA, the linear coefficient threshold and the amount of regularization were computed based on Bayesian optimization search. Finally, all ANN decoders consisted of one hidden layer with 128 nodes, followed by a sigmoid and a softmax layer, each with 4 nodes (for keypress decoding). Training utilized a scaled conjugate gradient optimizer with backpropagation, employing a learning rate of 0.01 (coarse to fine tuning) for a maximum of 100 epochs, with early stopping validation patience set to 6 epochs.

#### Decoding Performance Metric

Decoding performance was assessed using several metrics, including accuracy (%), which indicates the proportion of correct classifications among all test samples. Confusion matrices provide a detailed comparison of the number of correctly predicted samples for each class against the ground truth. The F1 score—defined as the harmonic mean of the precision (percentage of true predictions that are actually true positive) and recall (percentage of true positives that were correctly predicted as true) scores—was used as a comprehensive metric for each one-versus-all keypress state decoder to assess class-wise performance that accounts for both false-positive and false-negative prediction tendencies [79, 80]. A weighted mean F1 score was then computed across all classes to assess the overall prediction performance of the multi-class model. Test accuracies based on the best decoder performance (LDA in our case) were reported and used for statistical comparisons.

#### Decoding During Skill Learning Progression

We systematically assessed decoding performance of a 2-class decoder (Index_OP1_ vs Index_OP5_; i.e. – decoding of two index finger keypresses occurring at different locations within the training sequence) at each trial during the skill learning process to capture the evolving relationship between differentiated index finger decoding proficiency and the acquired skill. Our approach involved evaluating decoder performance individually for each *Day 1 Training* trial. We ensured an equal number of samples (first *k* keypresses) in each trial were used to mitigate the influence of increasing samples available in later trials.

We used t-distributed stochastic neighborhood estimation (t-SNE) to visualize the evolution of neural representations corresponding to each keypress at each trial of the learning period. Within t-SNE distributions, index finger keypresses were separately labeled based upon their sequence location (i.e., Index_OP1_ and Index_OP5_ for ordinal positions 1 and 5, respectively).

#### Decoding Sequence Elements

We performed 5-class decoding of each action based on its location within the sequence (i.e., Index_OP1_, Little, Middle, Ring, and Index_OP5_). The same decoding strategy was utilized as for the above 4-class keypress-based decoding (i.e. -90%-10% split for train and test, 8-fold cross validation of training samples to select best decoder configuration, hybrid spatial features, and LDA based dimension reduction). Note, oversampling was not needed after sub-grouping the index finger keypresses into two separate classes based on their sequence context. 5-class sequence-based decoding was evaluated for both *Day 1 Training*, *Day 2 Retest* and *Day 2 Control* data.

#### Feature Importance Scores

The relative contribution of source-space voxels and parcels to decoding performance (i.e. – feature importance score) was calculated using minimum redundant maximum relevance (MRMR) [81] and highlighted in topography plots. MRMR, an approach that combines both relevance and redundancy metrics, ranked individual features based upon their significance to the target variable (i.e. – keypress state identity) prediction accuracy and their non-redundancy with other features.

### Neural Representation analysis

We evaluated the *online* (within-trial) and *offline* (between-trial) changes in the neural representation of the contextual actions (Index_OP1_ and Index_OP5_) for each trial during training. For offline differentiation, we evaluated the Euclidian distance between the hybrid spatial features of the last index finger keypress of a trial (Index_OP5_) to the first index finger keypress (Index_OP1_) of the subsequent trial, mirroring the approach used to calculate micro-offline gains in skill. This offline distance provided insight into the net change in contextual representation of the index finger keypress over each interleaved rest break. For online differentiation, we calculated either the mean Euclidian distance between Index_OP1_ and Index_OP5_ of all the correctly typed sequences (*sequence-based*) or the distance between the first Index_OP1_ and last Index_OP5_ (*trial-based*) within the same practice trial. Online differentiation informed on the net change in the contextual representation of the index finger keypress occurring within each practice trial. Cumulative offline and online representation distances across participants were statistically compared using paired *t*-tests. As a control analysis, we computed the difference in neural representation between Index_OP1_ and Index_OP5_ on *Day 2 Retest* data for the same sequence (**4**-1-3-2-**4**) as well as for different *Day 2 Control* untrained sequences where the same action was performed at ordinal positions 1 and 5 (**2**-1-3-4-**2**; **1**-4-2-3-**1**; **2**-3-1-4-**2**; **4**-2-3-1-**4**). We also assessed for specificity of contextualization to the trained sequence, by evaluating differentiation between index finger keypress representations performed at two different positions within untrained sequences (**4**-2-**4**-3-1 and 1-**4**-3-**4**-2). The cumulative differences were compared across participants with paired *t*-tests.

Finally, we computed trial-by-trial differences in offline and online representations during early learning, exploring their temporal relationships with cumulative micro-offline and -online gains in skill, respectively, through regression analysis and Pearson correlation analysis. Linear regression models were trained utilizing the *fitlm* function in MATLAB. The model employed M-estimation, formulating estimating equations and solving them through the Iteratively Reweighted Least Squares (IRLS) method [82]. Key metrics such as the square root of the mean squared error (RMSE), which estimates the standard deviation of the prediction error distribution, the coefficient of explained variance (*R^2^*), the F-statistic as a test statistic for the F-test on the regression model, examining whether the model significantly outperforms a degenerate model consisting only of a constant term, and the *p*-value for the F-test on the model were computed and compared across different models. This multifaceted approach aimed to uncover the nuanced dynamics of neural representation changes in response to skill acquisition.

## Data Availability

All de-identified and permanently unlinked from all personal identifiable information (PII) data are publicly available (https://nih.box.com/v/hcpsSkillLearningData).

## Acknowledgments

We thank Ms. Tasneem Malik, Ms. Michele Richman, NIMH MEG Core Facility staff, and NIH NMRF and FMRIF Core Facility staff for their support. This work utilized the computational resources of the NIH HPC Biowulf cluster (http://hpc.nih.gov). This research was supported by the Intramural Research Program of the NINDS, NIH.

## Supplementary Materials

**Figure 1 - figure supplement 1:**
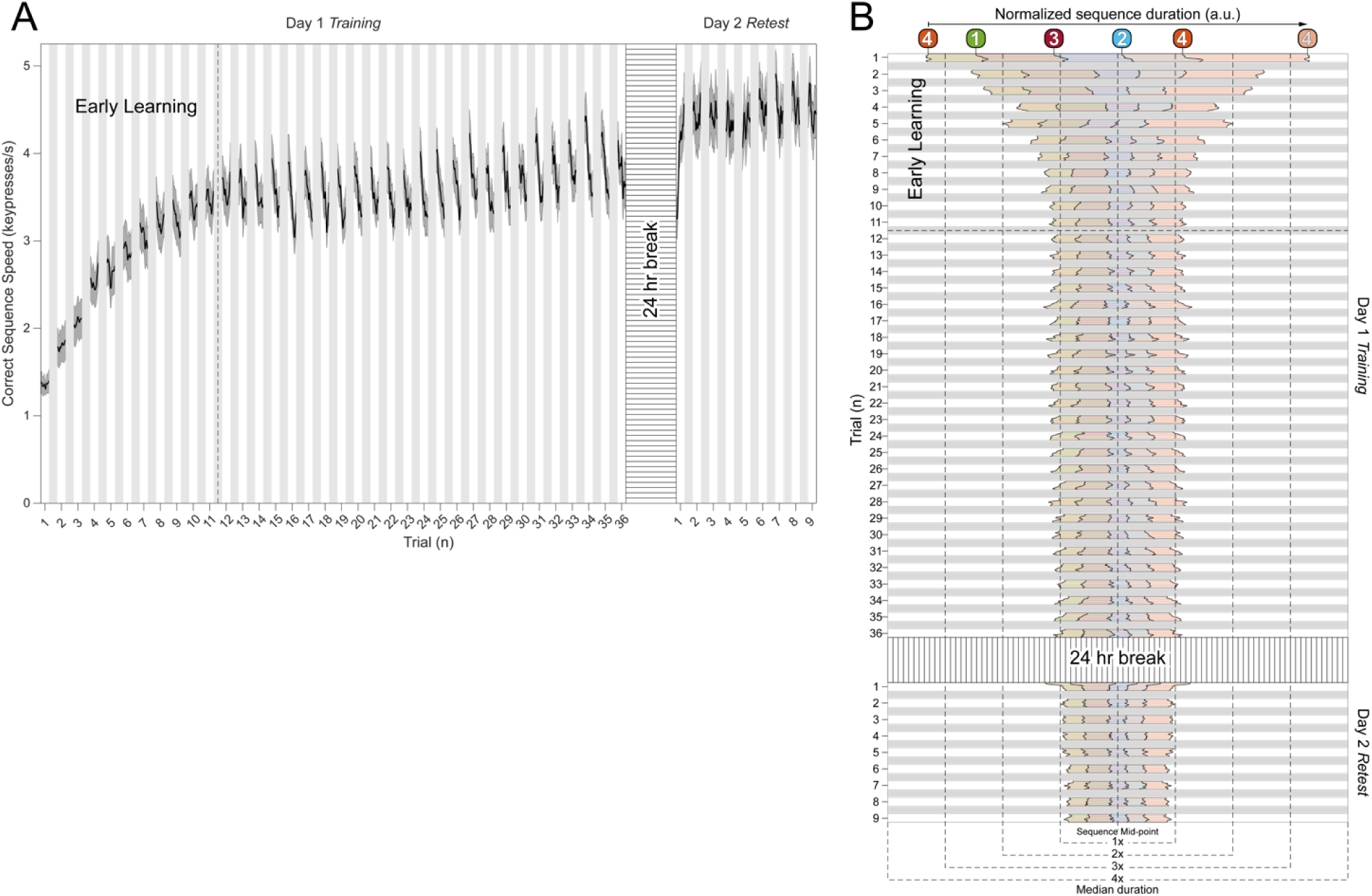
Behavioral performance during skill learning. A) Total Skill Learning over Day 1 Training (36 trials) and Day 2 Retest (9 trials). As reported previously [1], participants on average reached 95% of peak performance during *Day 1 Training* by trial 11. Note that after trial 11, performance stabilizes around a plateau through trial 36. Following a 24-hour break, participants displayed an upward shift in performance during the *Day 2 Retest* – indicative of an overnight skill consolidation effect. **B) Keypress transition time (KTT) variability.** Distribution of KTTs normalized to the median correct sequence time for each participant and centered on the mid-point for each full sequence iteration during early learning. Note that the initial variability of the five component transitions in the sequence (i.e. – 4-1, 1-3, 3-2, 2-4, 4-4) stabilize by trial 6 in the early learning period and remain stable throughout the rest of *Day 1 Training* (through trial 36) and *Day 2 Retest*.

**Figure 2 - figure supplement 1:**
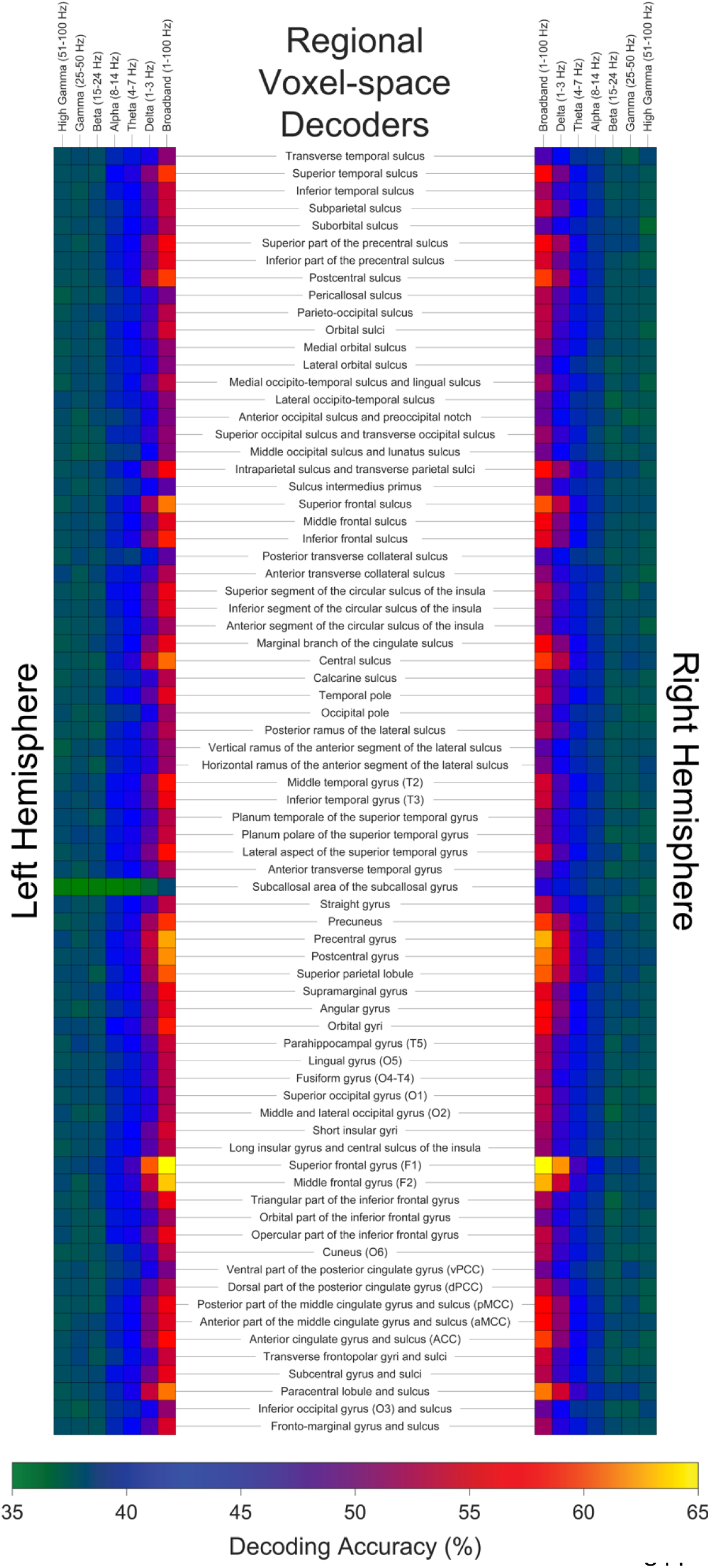
Oscillatory contributions at individual brain regions. Decoding performance of regional voxel-space activity patterns within individual brain areas for broadband and each narrowband oscillatory range is displayed in the form of a heatmap for both the left and right hemisphere. Optimal decoding performance for broadband regional voxel-space decoders were obtained from bilateral superior frontal (Left: 68.77% ± SD 7.6%; Right: 67.52% % ± SD 6.78%), middle frontal (Left: 63.41% ± SD 7.58%; Right: 62.78% % ± SD 76.94%), pre-central (Left: 62.37% % ± SD 6.32%; Right: 62.69% ± SD 5.94%), and post-central (Left: 61.71% ± SD 6.62%; Right: 61.09% ± SD 6.2%) brain regions. Superior parietal, central, paracentral, anterior-cingulate, and precuneus regions also showed broadband decoding performance exceeding 60%. With respect to decoders constructed from narrowband oscillatory input features, only Delta-band voxel-space activity from bilateral superior frontal regions achieved at least 60% decoding accuracy of keypresses.

**Figure 2 - figure supplement 2:**
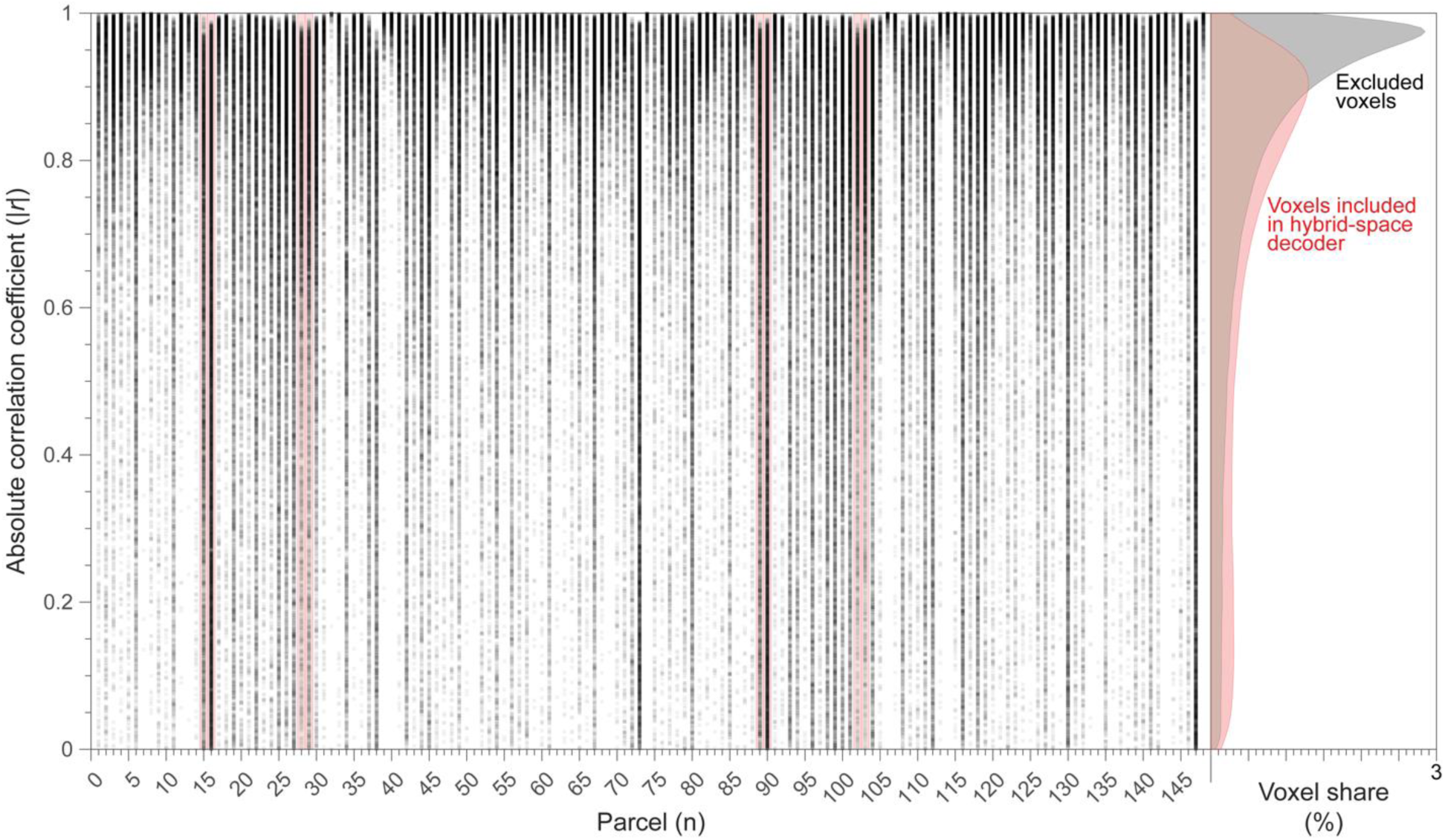
Distribution of correlation coefficients between parcel-space time-series and their constituent voxels. Data is shown for all subjects. Parcels represented in the regional voxel-space features of the hybrid-space decoder are marked with red vertical boxes (bilateral superior frontal, middle frontal, pre-central and post-central regions). The y-axis indicates the absolute correlation coefficients for each voxel time-series with the time-series of the parcel it is a member of (1 = complete redundancy; 0 = orthogonality). Note that while signal in some voxels correlate strongly with parcel-space time-series, others are fully orthogonal. That is, the degree to which information obtained at the two different spatial scales is complimentary (or redundant) varies substantially over the regional voxel-space. This finding is consistent with the documented increase in correlational structure of neural activity across larger spatial scales that does not reflect perfect dependency or orthogonality [28]. The normalized cumulative distributions of parcel-to-voxel-space correlations depicted on the right show that voxels included in the hybrid-space decoder (red) are correlated less overall (two-sample Kolmogorov-Smirnov test: *D* = 0.2484, *p* < 1x10^-10^) with their respective parcel-space time-series relative to excluded voxels (grey).

**Figure 3 - figure supplement 1:**
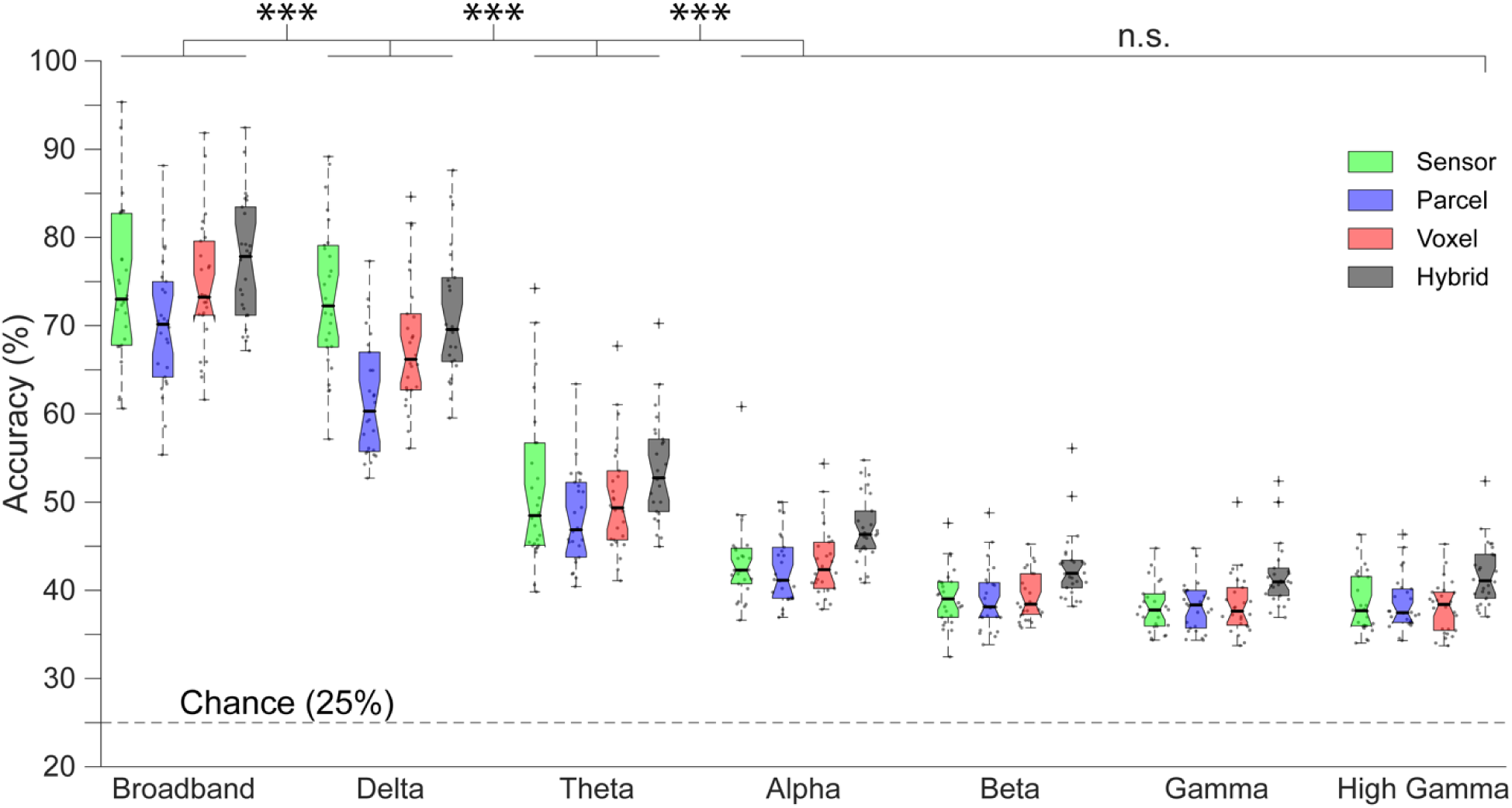
Contribution of whole-brain oscillatory frequencies to decoding. Accuracy for decoders trained on four different input feature spaces—sensor, whole-brain parcel, whole-brain voxel and hybrid (combination of whole-brain parcel plus regional voxel)—was highest for broadband MEG activity, followed by Delta-band activity. The hybrid approach resulted in the highest decoding accuracy, regardless of whether input features were broadband or narrowband-limited. Sensor-, parcel- and voxel-space decoders displayed similar accuracy with respect to one another for broadband MEG activity, and also for all narrowband ranges assessed. Dots depict decoding accuracy for each participant. “**\*\*\***” indicates 𝑝 < 0.001, while “n.s.” denotes no statistical significance (i.e. - 𝑝 > 0.05).

**Figure 3 - figure supplement 2:**
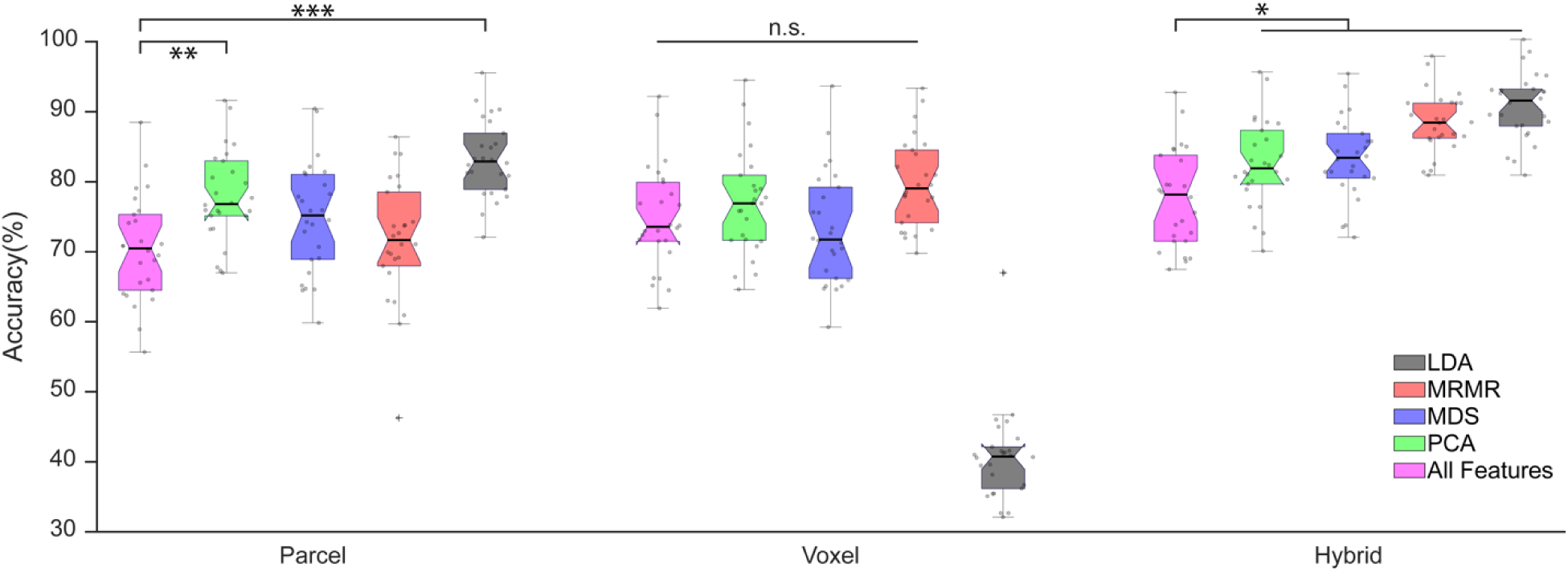
Comparison of different dimensionality reduction techniques. Dimensionality reduction was applied to the input features for each approach (parcel-space: N=148; voxel-space: N=15684; hybrid-space: N=1295)[34]. The results with principal component analysis (PCA, in green), multi-dimensional scaling (MDS, in blue), minimum redundant maximum relevance algorithm (MRMR, in red), linear discriminant analysis (LDA, in black) are shown in comparison to performance obtained using all input features (in magenta). For parcel-space input features, all these approaches increased the mean decoding accuracy with PCA and LDA (both of which result in extraction of orthogonal features) showing statistically significant improvement (1-way ANOVA: F= 13.05, *p* < 0.001; post hoc Tukey tests: *p* =0.032; PCA: *p* < 0.001; LDA: *p* > 0.05). For voxel-space features, there was no statistically significant improvement with any of the approaches (*p* > 0.05). While MRMR resulted in the largest voxel-space decoding accuracy improvement it was not statistically significant (post hoc Tukey test: *p* = 0.14), and application of LDA dimensionality reduction actually reduced performance dramatically. Uniquely for hybrid-space features—all dimensionality reduction techniques improved decoding performance significantly (1-way ANOVA: F= 21.32; post hoc Tukey tests: *p* < 0.05) with the best largest improvement observed following application of LDA. “**\*\*\***” indicates 𝑝 < 0.001, “**\*\***” indicates 𝑝 < 0.01, “**\***” indicates 𝑝 < 0.05 and “n.s.” denotes no statistical significance (i.e. - 𝑝 > 0.05).

**Figure 3 - figure supplement 3:**
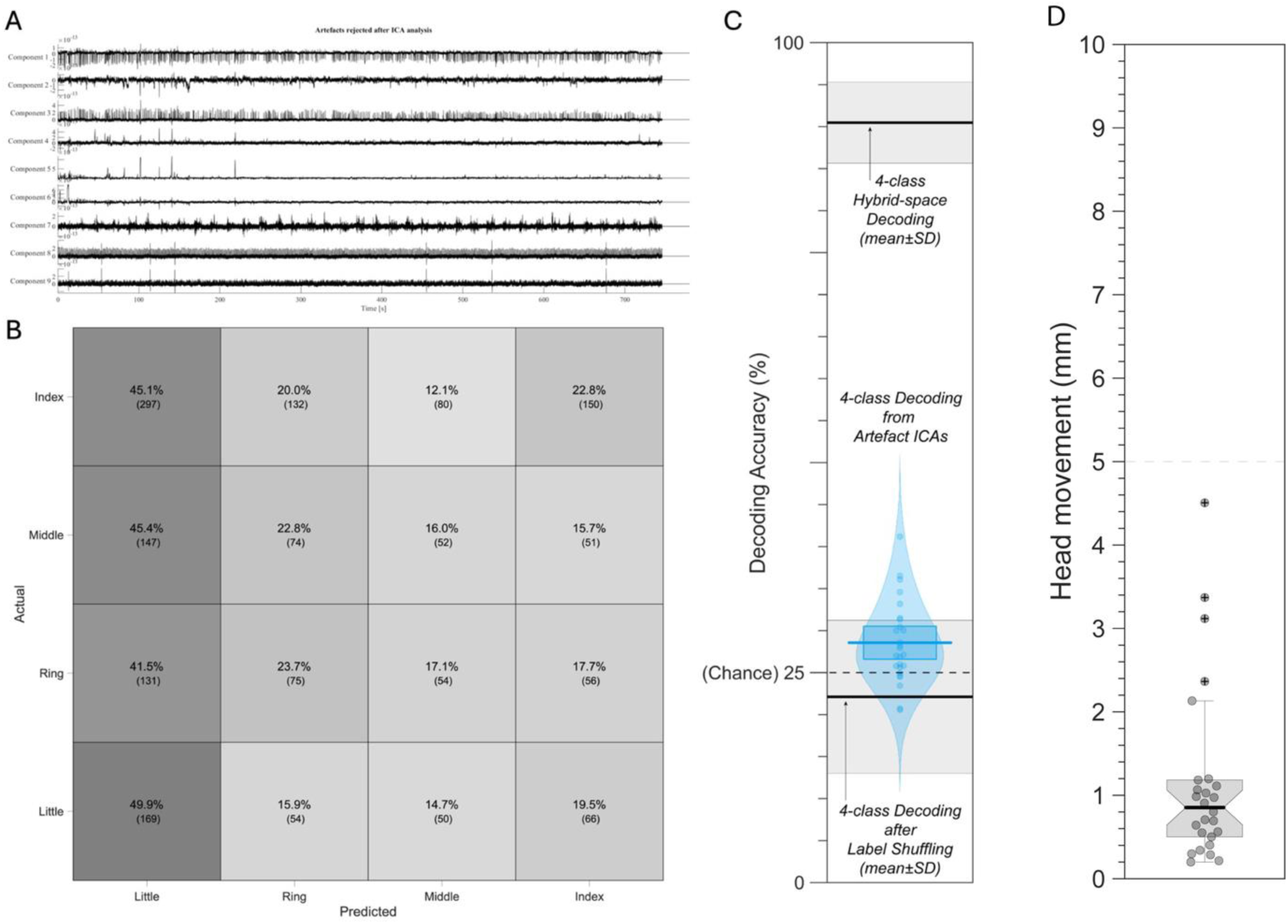
**A)** Example of ICA component time-series for components labeled as artefacts from a single subject during MEG data pre-processing. The features of these components are consistent with known motion and physiological artefacts in MEG data. **B)** 4-class confusion matrix and **C)** decoding performance of keypress action labels from ICA components labeled as artefacts and removed from the MEG data during pre-processing. These components failed to predict keypress labels above empirically determined chance levels (as shown by decoding performance after random label shuffling). Note that in all cases, decoding performance from movement and physiological artefacts was substantially lower than 4-class MEG hybrid-space decoding for all participants. **D)** Head position was assessed at the beginning and at the end of each recording and used to measure head movement. The mean measured head movement across the study group was 1.159 mm (± 1.077 SD).

**Figure 3 - figure supplement 4:**
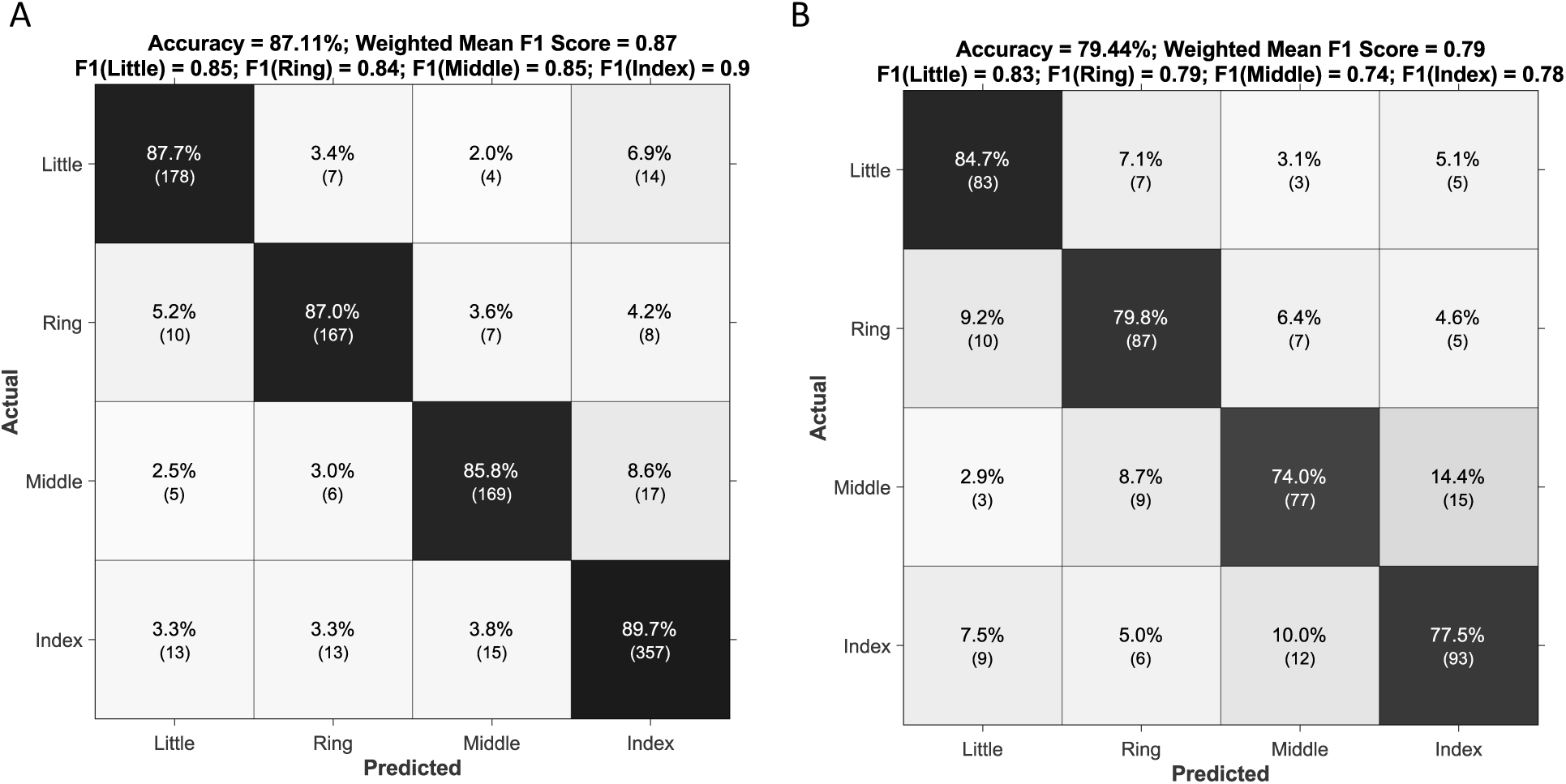
Confusion matrices for decoding performance on *Day 2 Retest* **(A)** and *Day 2 Control* **(B)** data. Note that the hybrid-space decoding strategy generalized to Day 2 data with 87.11% overall accuracy for keypresses embedded within the trained sequence (*Day 2 Retest*) and 79.44% overall accuracy for keypresses embedded within untrained control sequences (*Day 2 Control*).

**Figure 3 - figure supplement 5:**
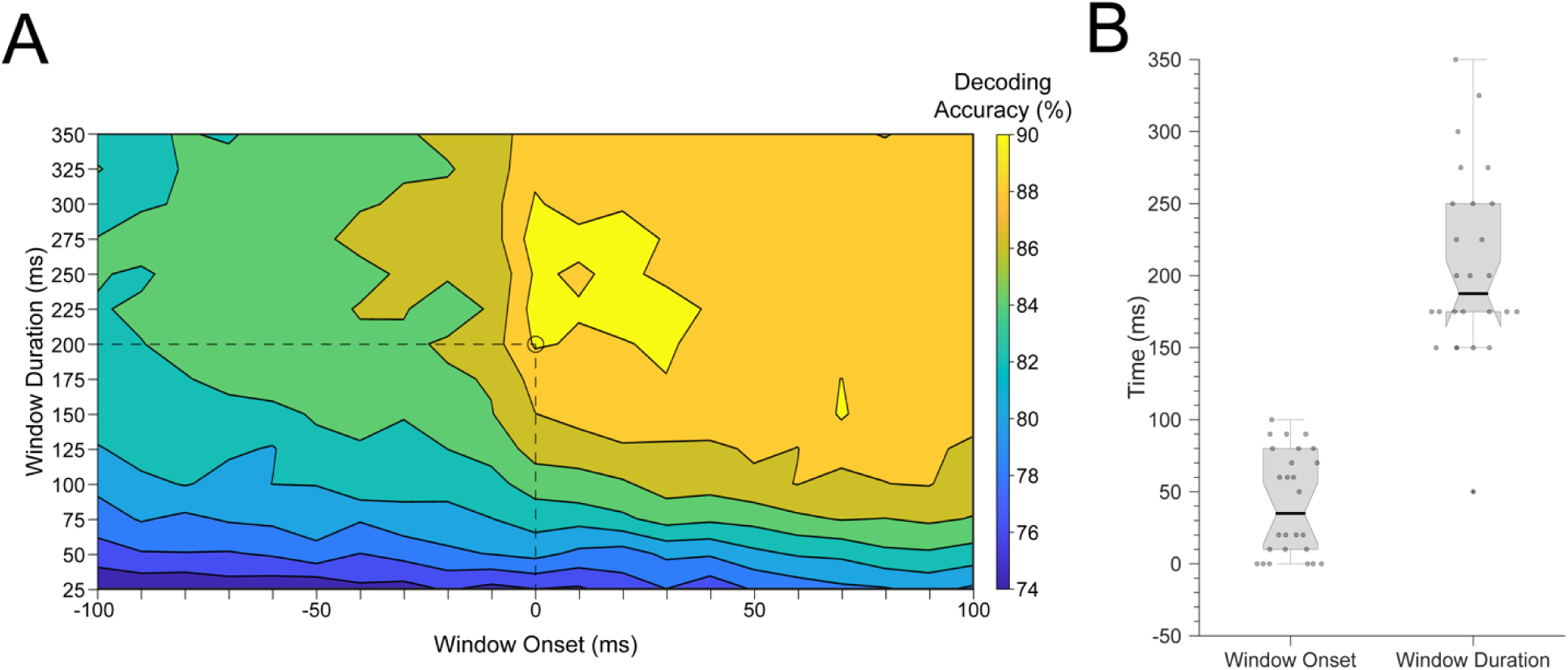
**A)** Average decoding accuracies across participants with varying window parameters. The x-axis indicates the onset of the time window (in ms) used to relate MEG activity time-series to individual keypresses (i.e. – KeyDown event = 0 ms), while the y-axis indicates the window duration (in ms). The heatmap color denotes the decoding accuracy for all window onset/duration pairings. The best decoding accuracy across subjects was obtained using a window duration of 200 ms with the leading edge aligned to the KeyDown event (i.e. – 0 ms; marked by the dashed lines and open circle). **B)** Decoder window parameters (onset and duration) used for each subject in reported decoder accuracy comparisons (Figures 2-4). Please note that the group-optimal set of parameters (window onset = 0 ms; window duration = 200 ms; LDA dimensionality reduction) was utilized for all contextualization analyses (Figure 5) to allow for comparison across participants.

**Figure 3 - figure supplement 6:**
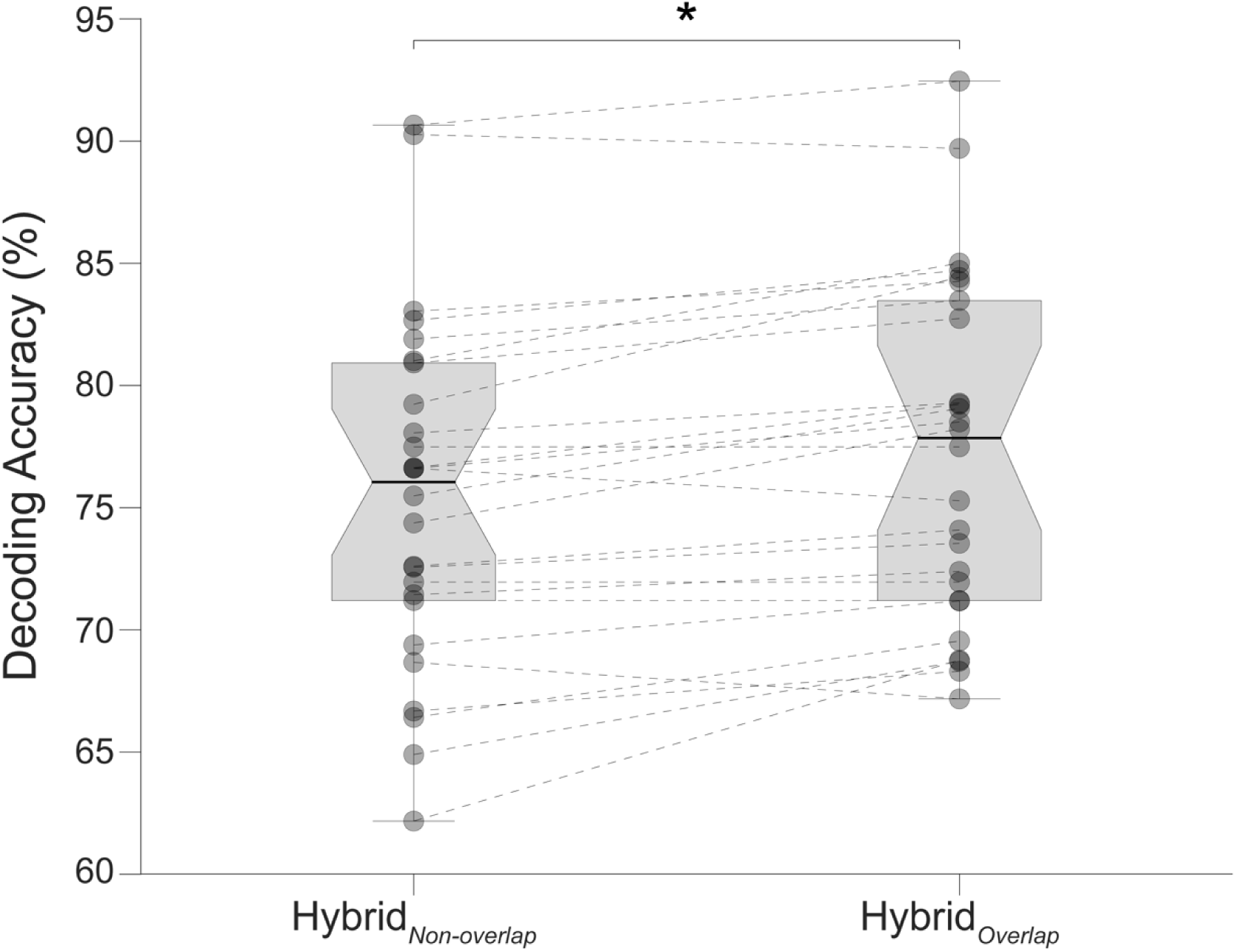
Comparison of decoding performances with two different hybrid approaches. Hybrid_Overlap_ (regional voxel-space features from top-ranked parcels combined with all whole-brain parcel-space features as shown in Figures 3B and **Figure 3 – figure supplements 1,3-5** of the manuscript) and Hybrid_Non-overlap_ (regional voxel-space features of top-ranked parcels and spatially *non-overlapping* whole-brain parcel-space features). Filled circle markers represent decoding accuracy for individual subjects. Dashed lines indicate within-subject performance changes between decoding approaches. Note, that the Hybrid_Overlap_ (the approach used in our manuscript) significantly outperforms the Hybrid_Non-overlap_ approach (Wilcoxon signed rank test, *z* = 3.7410, *p* = 1.8326e-04), despite the removed features (n = 8) only comprising less than 1% of the overall input feature space. These results indicate that the spatially overlapping *whole-brain* (lower resolution) *parcel-space* and *regional* (higher resolution) *voxel-space* features provide complimentary—as opposed to redundant—information to the hybrid-space decoder.

**Figure 3 - figure supplement 7:**
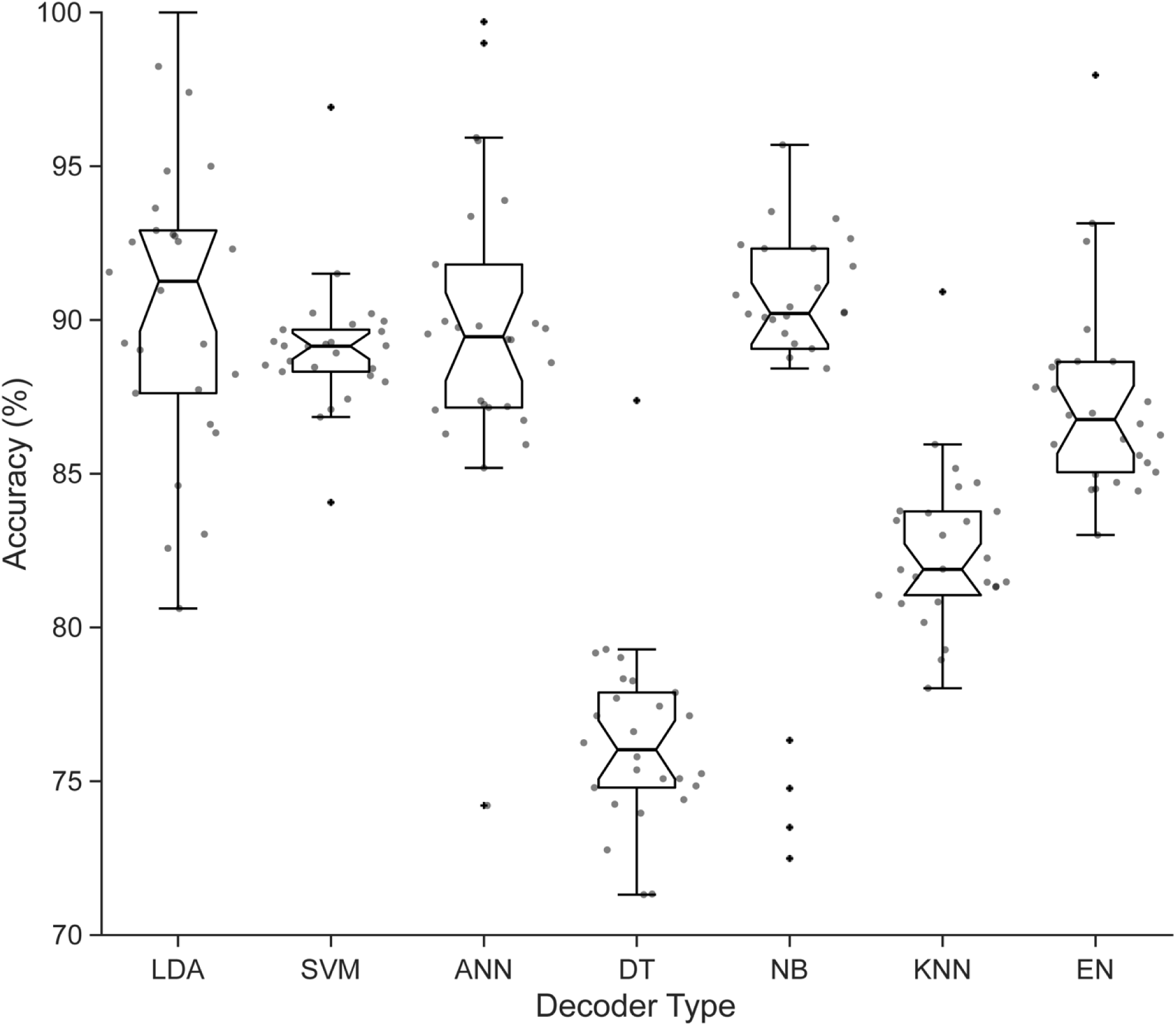
Comparison of different decoder methods. Performance for all different machine learning decoders assessed is shown for each participant. The results show that the linear discriminant analysis (LDA) classifier outperformed other methods, on average, across the group. Decoding analysis performance comparisons reported in the current study utilized the LDA decoder for all subjects.

**Figure 4 - figure supplement 1:**
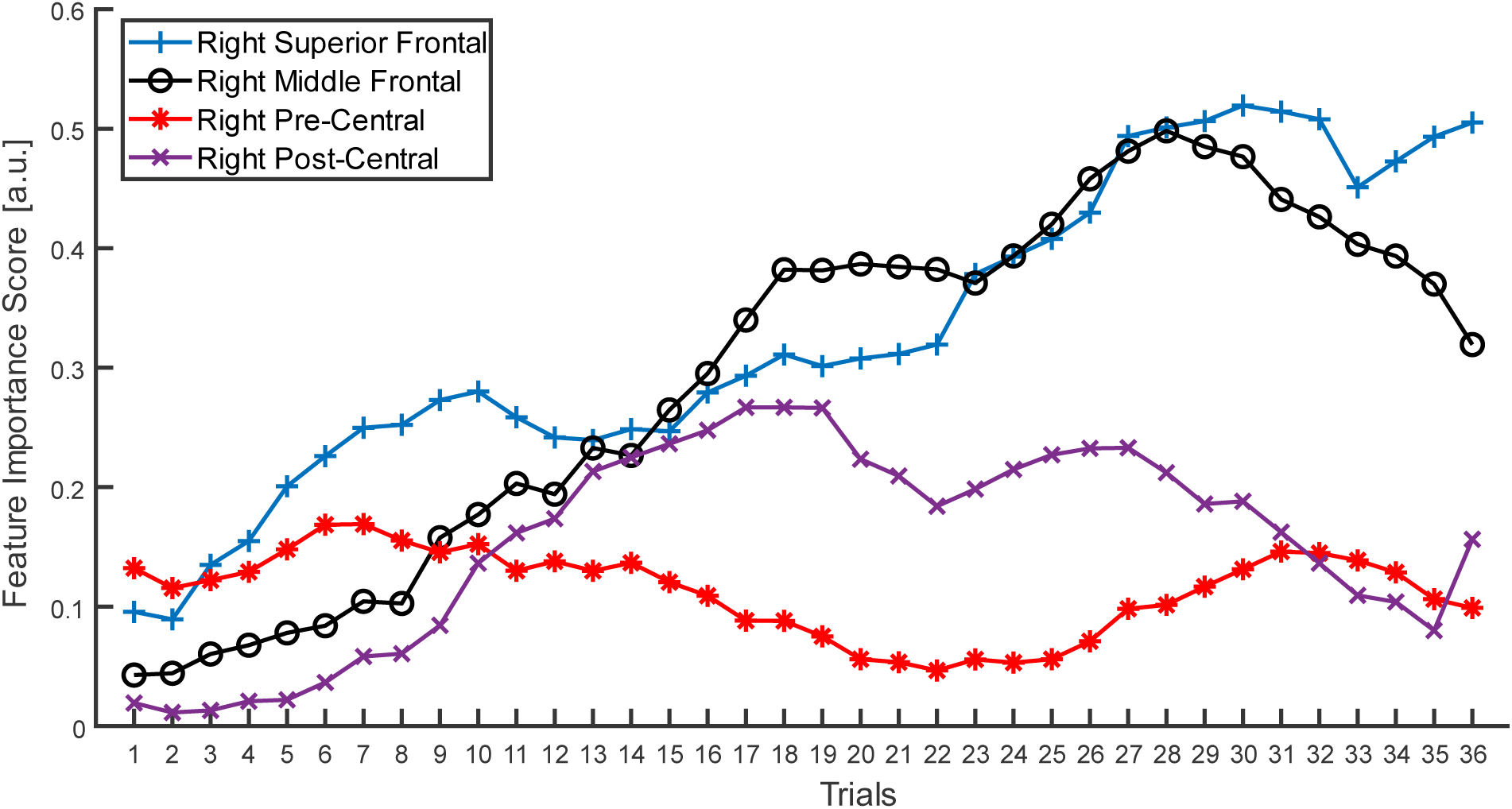
Quantification of parcel-space trial-by-trial feature importance score during skill learning. Parcel-space trial-by-trial changes in feature importance scores are shown for right superior frontal, middle frontal, pre-central, and post-central cortex (i.e. – the contralateral regions showing the highest regional voxel-space decoding accuracy). Note that the feature importance is initially higher for the contralateral pre-central cortex in early trials before shifting towards the contralateral middle and superior frontal cortex during later trials, as can be seen with the divergence of line plots beginning around trial 11.

**Figure 4 - figure supplement 2:**
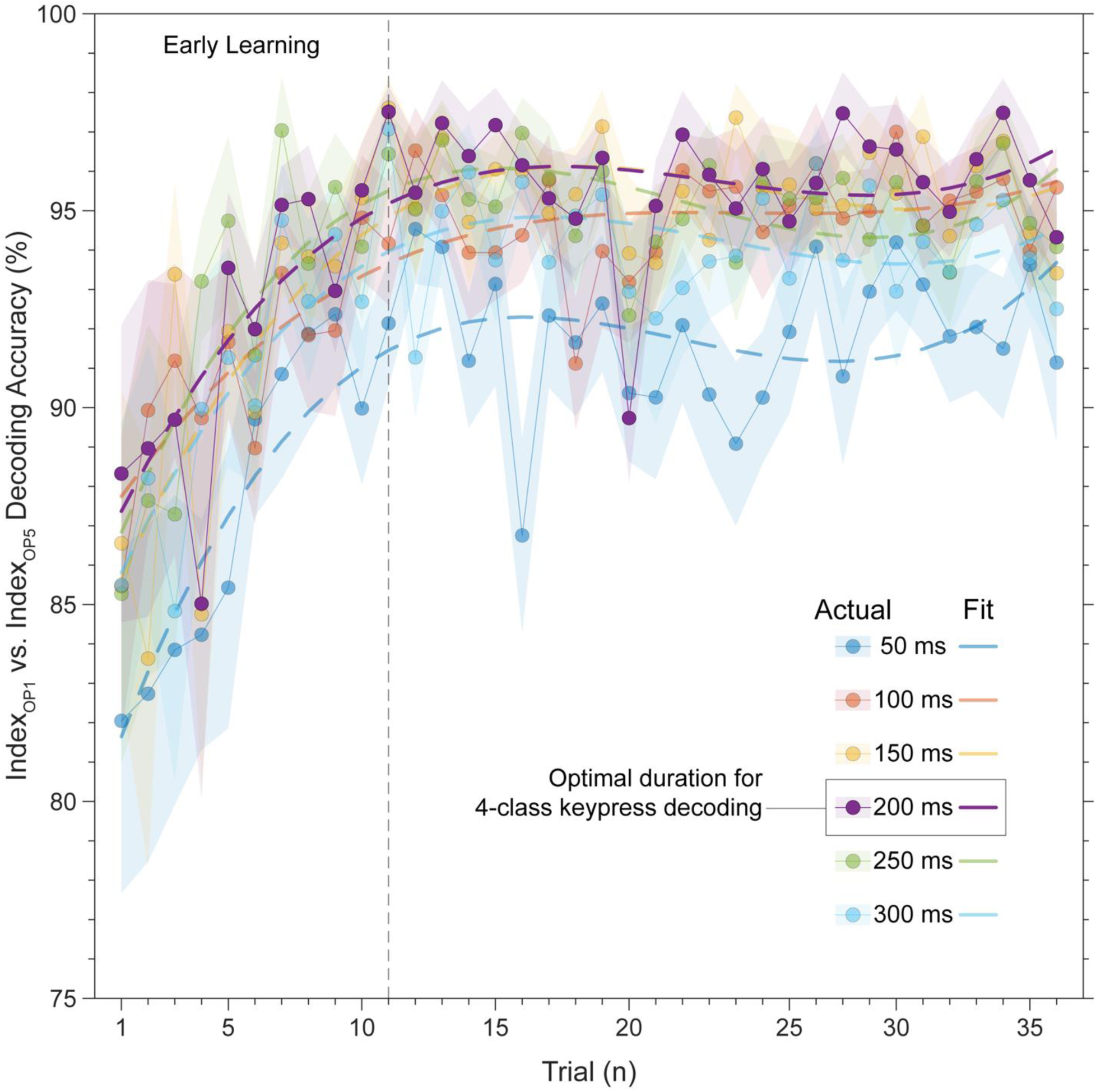
Trial-by-trial classification accuracy for 2-class decoder (Index_OP1_ vs. Index_OP5_). Several decoders (with varying window durations aligned to the KeyDown event) were trained to differentiate between the two index finger keypresses embedded at different positions within the practiced skill sequence (Index_OP1_ at ordinal position 1 vs. Index_OP5_ at ordinal position 5). Decoding accuracy for the 200 ms duration windows (i.e. – the optimal window size for 5-class decoding of individual keypresses) progressively improves over early learning, stabilizing around 96% by trial 11 (end of early learning). Similar results were observed for all other decoding window sizes (50, 100, 150, 250 and 300 ms), with overall accuracy slightly lower compared to 200 ms. These findings indicate that the neural representations of the skill action is updated over early learning to incorporate sequence location information.

**Figure 4 - figure supplement 3:**
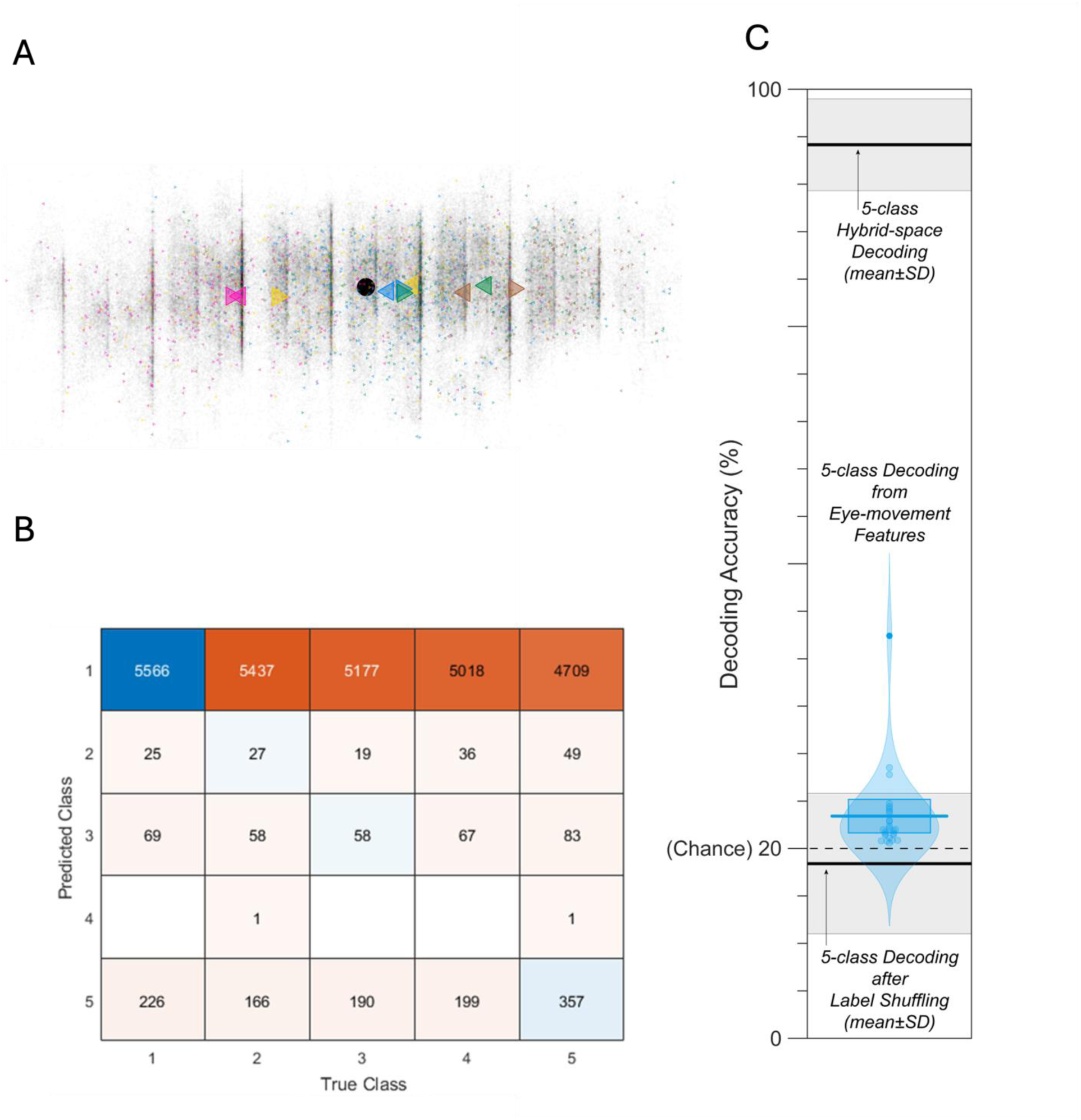
**A)** Scatter plot of gaze positions at the KeyDown event and 200ms after the KeyDown event (i.e. – beginning and ending of window used for decoding keypress labels from MEG input features) from a representative participant. Transparent grey dots indicate all sampled gaze positions during practice trials. The overall mean gaze position during practice trials is indicated by the black filled circle marker. Colored right-pointing triangle markers indicate the gaze position at the KeyDown event for each ordinal position keypress (Index_OP1_ – magenta; Little_OP2_ – yellow; Middle_OP3_ – blue; Ring_OP4_ – green; Index_OP5_ – brown), while left-pointing triangle markers indicate the gaze position 200ms after the KeyDown event. The mean gaze position for these two time-points is indicated by the larger-sized triangle markers. On average, gaze position is largely fixed for the OP1 and OP3 keypresses, moves from left-to-right for OP2 and OP4 keypresses, and from right-to-left for OP5 keypresses (which is when the asterisk moves leftward from the last sequence item back to the first). **B)** Confusion matrix showing that three eye movement features fail to predict asterisk position on the task display above chance levels (Fold 1 test accuracy = 0.21718; Fold 2 test accuracy = 0.22023; Fold 3 test accuracy = 0.21859; Fold 4 test accuracy = 0.22113; Fold 5 test accuracy = 0.21373; Overall cross-validated accuracy = 0.2181). Since the ordinal position of the asterisk on the display is highly correlated with the ordinal position of individual keypresses in the sequence, this analysis provides strong evidence that keypress decoding performance from MEG features is not explained by systematic relationships between finger movement behavior and eye movements (i.e. – behavioral artefacts). **C)** 5-class decoding of ordinal position keypress labels from eye movement recording features approached empirically determined chance levels (as shown by decoding performance after random label shuffling). Note that all decoding performances from eye movement data was substantially lower than MEG hybrid-space decoding for all participants.

**Figure 5 - figure supplement 1.**
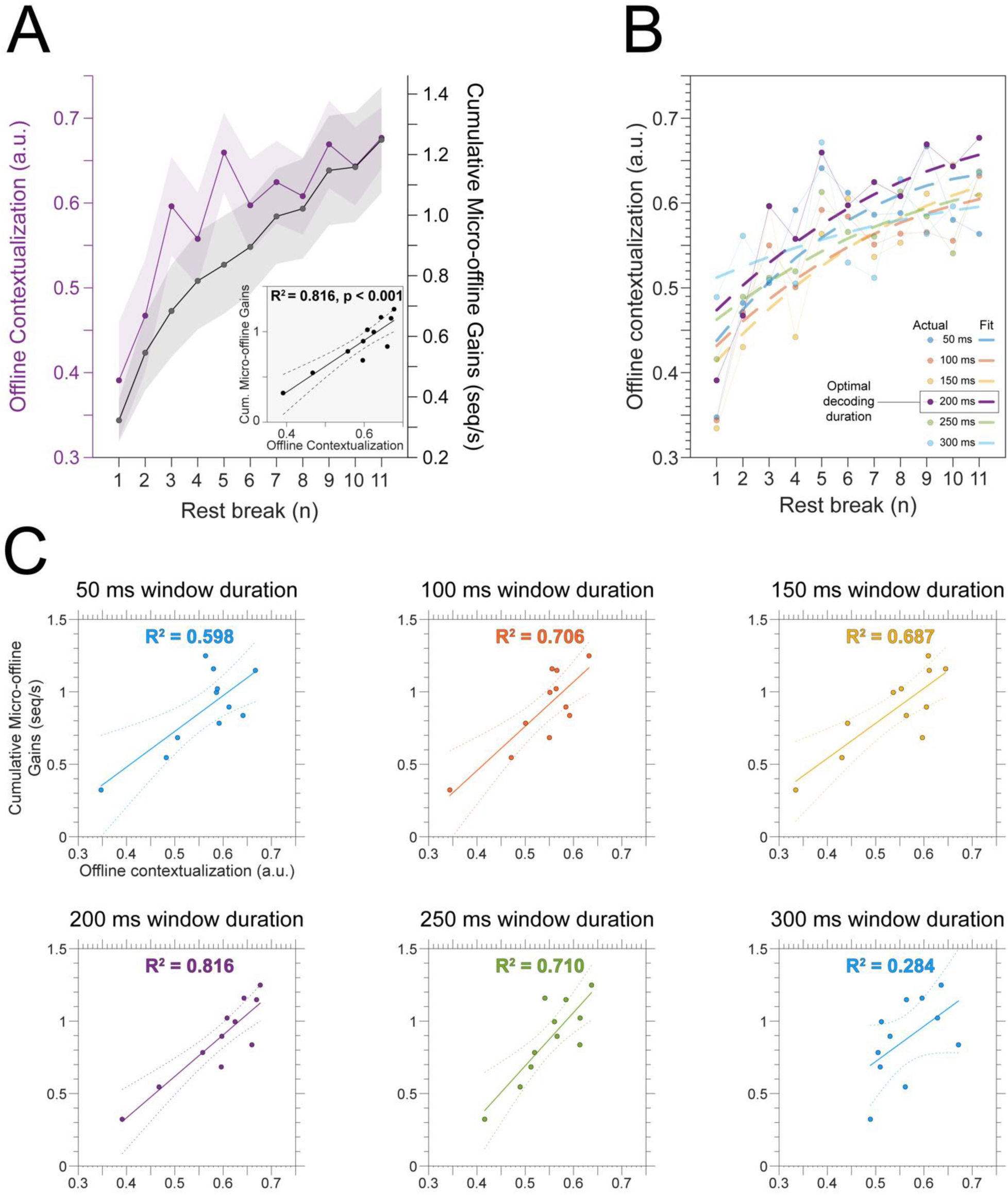
A) Relationship between offline neuronal representational changes and micro-offline learning. Offline contextualization—calculated as the Euclidian distance between the neural representations observed for the first Index_OP1_ keypress from practice trial, *n*, and the last Index_OP5_ keypress from practice trial, *n-1*—increased over early learning. A linear regression analysis (shown in the inset) revealed a strong temporal relationship (correlation coefficient [*r*] = 0.903 and coefficient of variance explained [R^2^] = 0.816) between contextualization and cumulative micro-offline gains over early learning. **B) Changes in offline contextualization for different decoding window durations as a function of rest breaks.** We constructed decoders from different MEG input feature time windows (window durations of 50, 100, 150, 200, 250 and 300ms; all aligned to the KeyDown event), to assess the robustness of the offline contextualization finding with respect to this parameter selection. Offline contextualization showed similar trends for all options tested. **C) Relationship between offline neural representational changes and micro-offline learning across all window durations.** The linear regression analysis from **(A)** was repeated for all contextualization measures from **(B)** obtained after varying the MEG input feature window size (50 – 300 ms). This strong temporal relationship was observed for all window durations (0.598 ≥ R^2^ ≥ 0.816), except for 300 ms (R^2^ = 0.284) where temporal overlap of individual keypress features was most prominent.

**Figure 5 - figure supplement 2:**
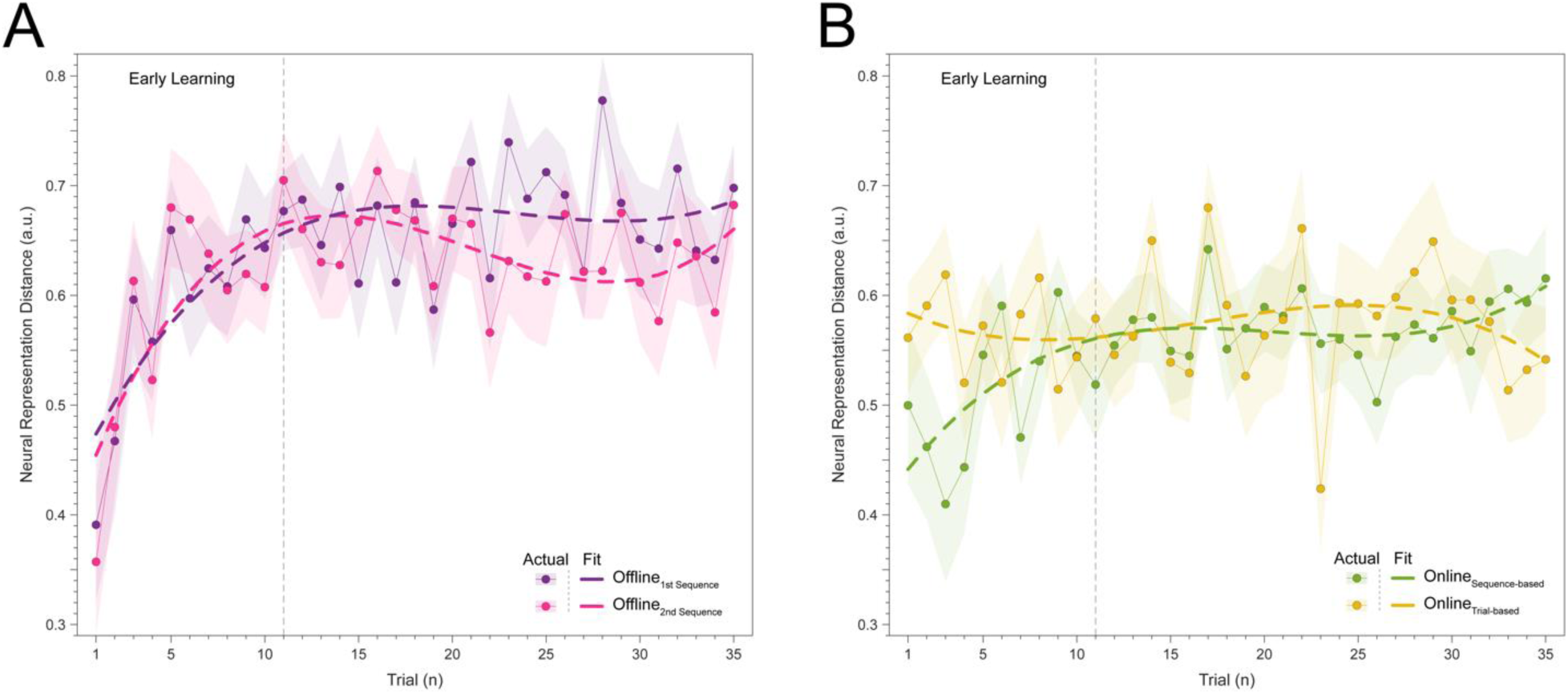
Trial-by-trial trends for different measurement approaches of offline and online contextualization changes. **A)** Offline contextualization between the last sequence of a preceding trial and the <u>second</u> sequence of the subsequent one (skipping the first sequence of that trial) rendered a comparable result to the measure reported in Figure 5 and **Figure 5 – figure supplement 1** which use the first sequence—inconsistent with a possible confounding effect of pre-planning [30]. **B)** Two different measurement approaches were used to characterize online contextualization changes. The *sequence-based* approach calculated the mean distance between Index_OP1_ and Index_OP5_ for each correct sequence iteration within a trial (green). A second *trial-based* approach was also implemented, which controlled for the passage of time between observations used in both online and offline distance measures (10 seconds between Index_OP1_ and Index_OP5_ observations in both cases). Note that the *trial-based* approach showed no increase in online contextualization over early learning. Importantly, the overall magnitude of online contextualization by the end of early learning was similar for both measurement approaches, and both showed reduced online relative to offline contextualization.

**Figure 5 - figure supplement 3:**
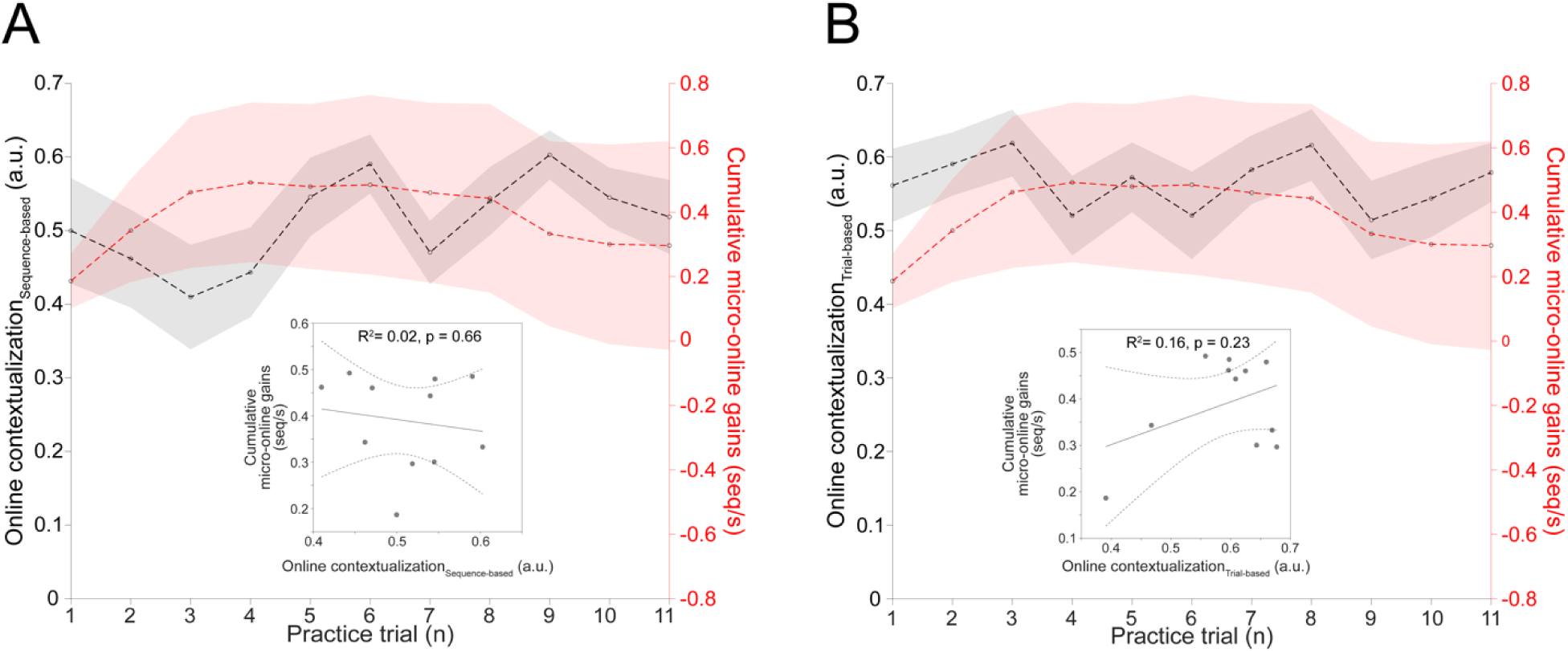
Online contextualization versus micro-online learning. The relationship between online contextualization and online learning is shown for both *sequence-* (**A**, left) and *trial-based* (**B**, right) distance measurement approaches. There was no significant relationship between online learning and online contextualization regardless of the measurement approach.

**Figure 5 - figure supplement 4:**
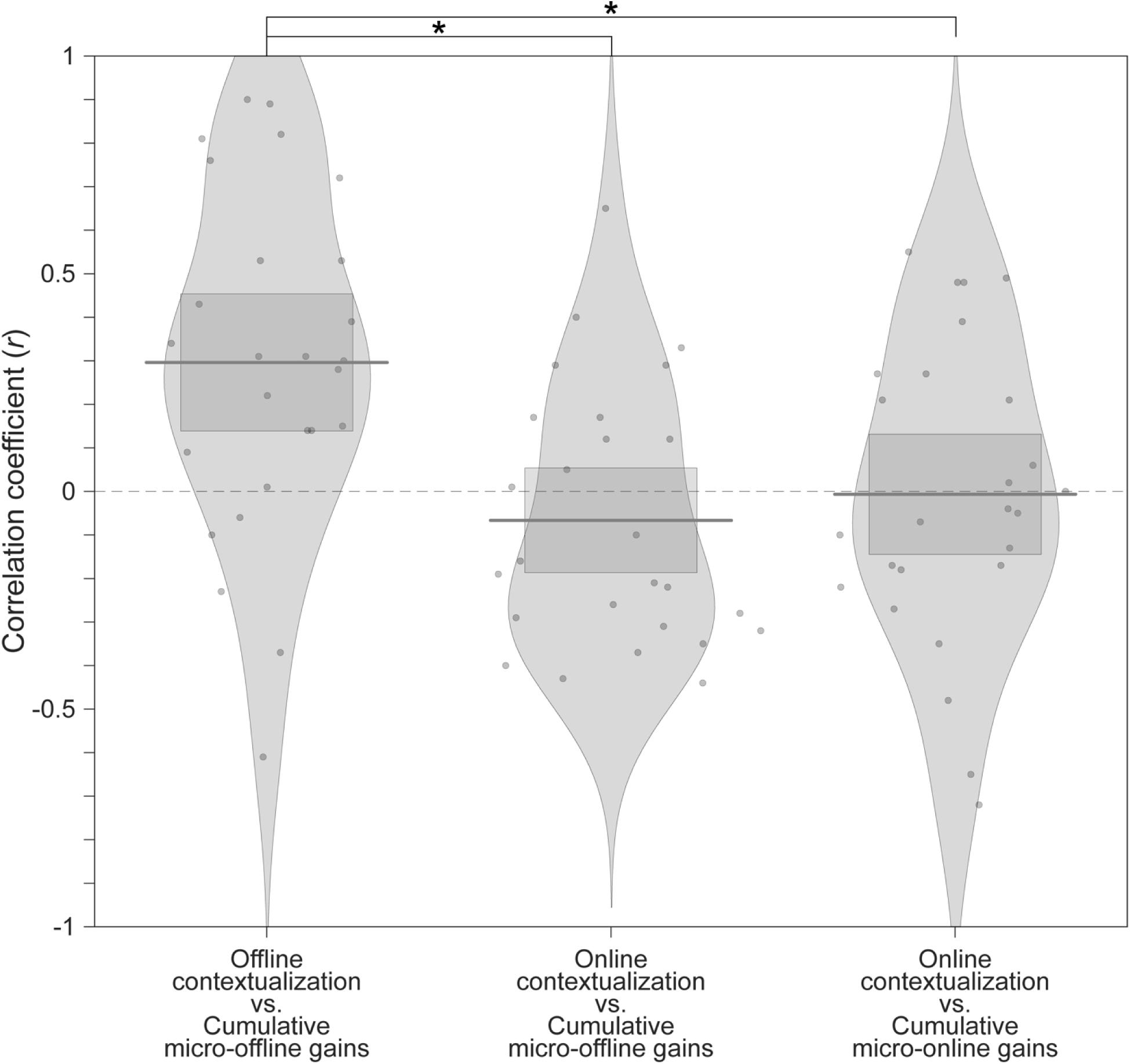
Within-subject correlations between online and offline contextualization changes versus learning. Pirate plots displaying individual subject correlation coefficients for offline (i.e. – over rest) and online (i.e. – during practice) contextualization changes versus micro-offline and -online performance gains. Within-subject correlations were significantly greater for offline contextualization changes versus micro-offline performance gains than for online contextualization changes versus either micro-offline or -online performance gains. The average correlation between offline contextualization and micro-offline gains within individuals was significantly greater than zero (**left;** *t* = 3.87, *p* = 0.00035, *df* = 25, Cohen’s d = 0.76) and stronger than correlations between online contextualization and either micro-online (**middle**; *t* = 3.28, *p* = 0.0015, *df* = 25, Cohen’s *d* = 1.2) or micro-offline gains (**right**; *t* = 3.7021, *p* = 5.3013e-04, *df* = 25, Cohen’s *d* = 0.69).

**Figure 5 - figure supplement 5:**
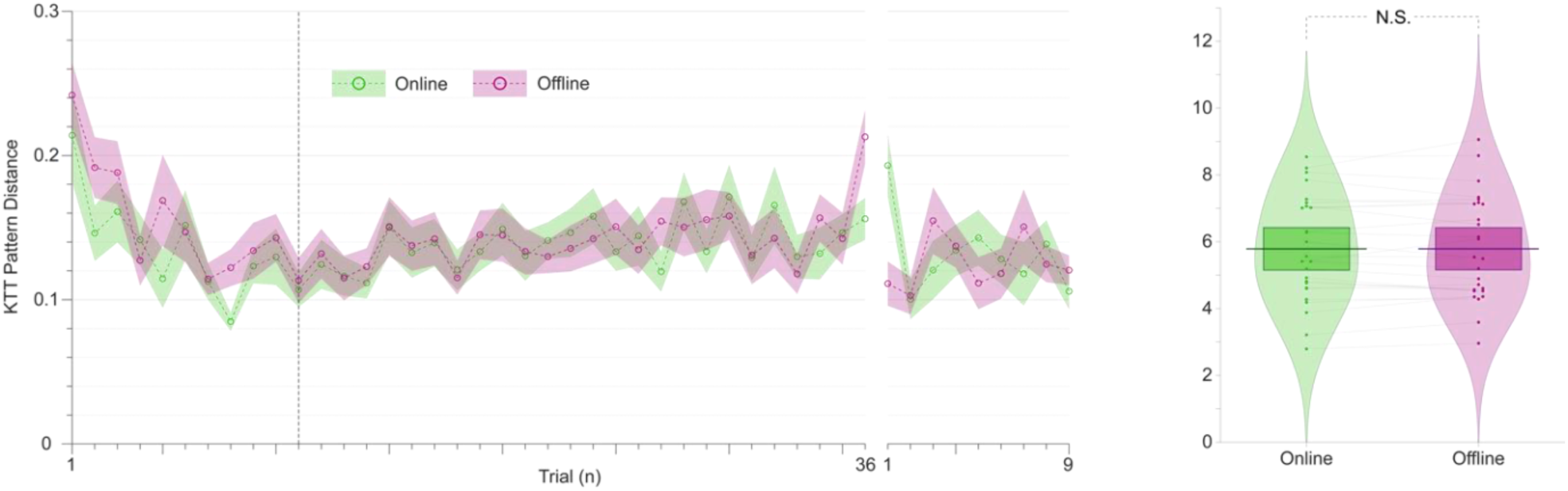
Online versus offline changes in keypress transition patterns. **A)** Trial-by-trial Euclidian distance between the relative share of each keypress transition time to the full sequence duration (i.e. – differences in typing rhythm). This distance was calculated for the first and last sequence of each trial (online pattern distance; green) and the last sequence of a trial versus the first sequence of the next (offline pattern distance; purple). **B)** Cumulative online (green; left) and offline (purple; right) pattern distances recorded over all forty-five trials covering Days 1 and 2. Note the comparable online and offline typing rhythm changes do not explain differences between online and offline contextualization, which is fully developed by trial 11 (Figure 5).

**Figure 5 - figure supplement 6:**
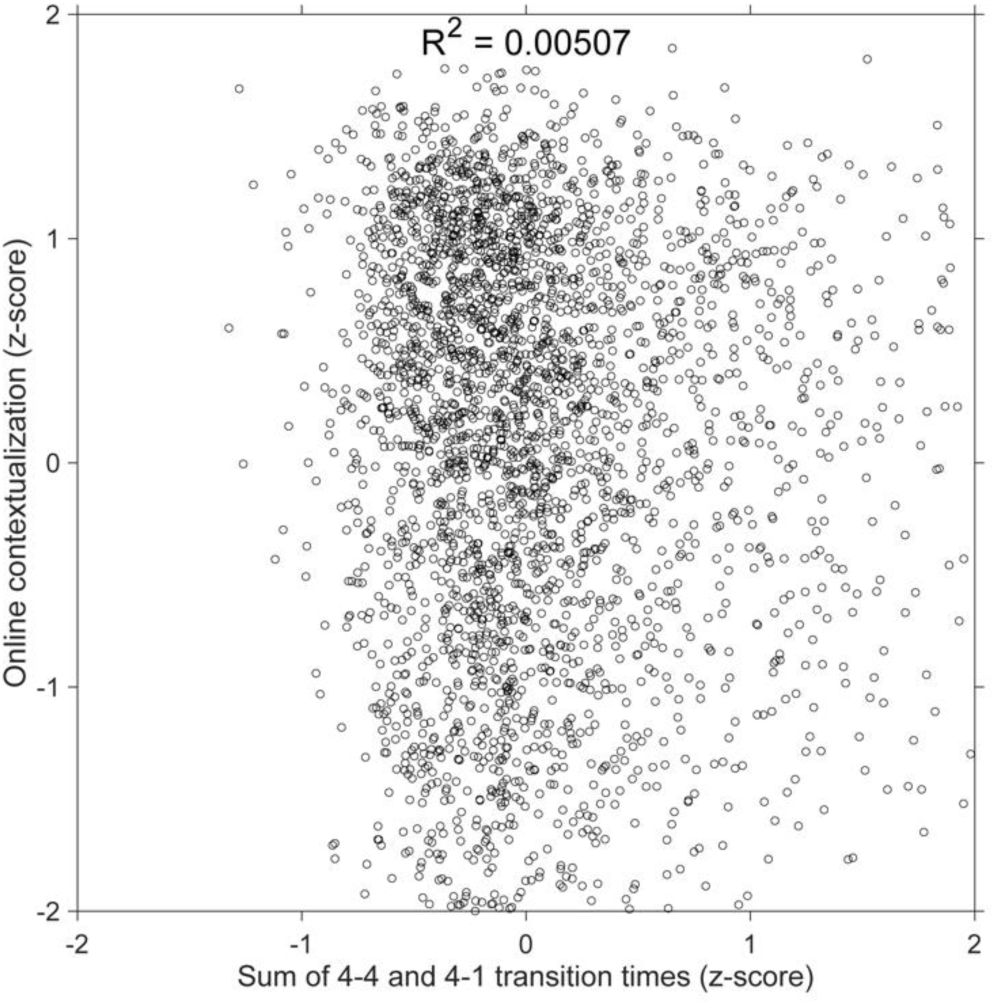
Relationship between adjacent index finger transitions and online contextualization. Scatter plot showing that the sum of adjacent index finger keypress transition times (i.e. – the 4-4 transition at the conclusion of one sequence iteration and the 4-1 transition at the beginning of the next sequence iteration) versus online contextualization distances measured during practice trials. Both the keypress transition times, and online contextualization scores were z-score normalized within individual subjects and then concatenated into a single data superset. A simple linear regression between keypress transition time predictor and the online contextualization response variable showed a very weak linear relationship between the two (R^2^ = 0.00507, F[1,3202] = 16.3). This result shows that contextualization of index finger representations does not reflect the amount of overlap between adjacent keypresses.

**Figure 5 - figure supplement 7:**
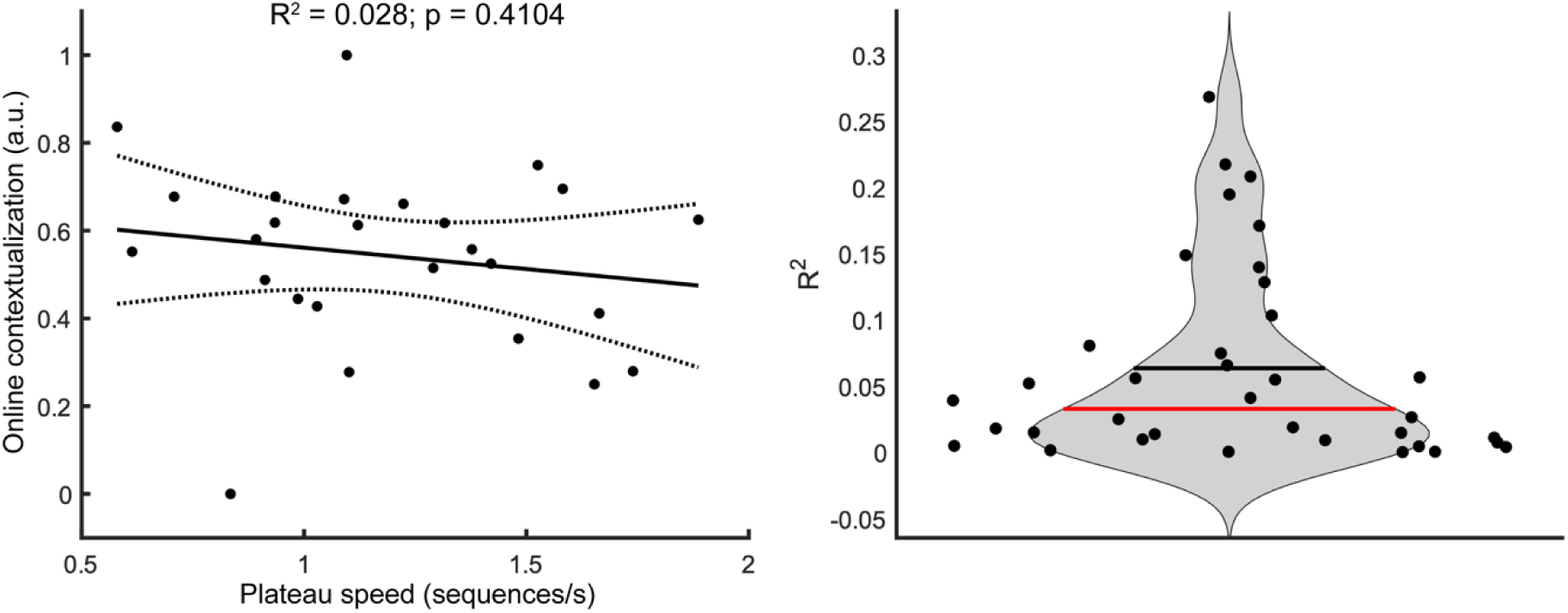
Between subject differences in typing speed versus online contextualization. **A)** Between-subject relationship between plateau performance speed and online contextualization. The plateau performance typing speed showed no significant relationship with the degree of online contextualization (*R^2^* = 0.028, *p* = 0.41). Each dot represents the maximum speed attained and the corresponding degree of contextualization of each participant. Thus, the magnitude of online contextualization was not dependent on how fast individuals could perform the task at the end of early learning. **B)** Trial-by-trial relationship between typing speed and degree of online contextualization. We also performed a trial-by-trial regression analysis that related the degree of online contextualization for each trial with the median typing speed for that trial. The R^2^ values obtained for regression analyses performed on individual trials were also low, and not statistically significant (mean *R^2^*= 0.06; *p* > 0.05).

